# The endoplasmic reticulum stress sensor IRE1 regulates collagen secretion through the enforcement of the proteostasis factor P4HB/PDIA1 contributing to liver damage and fibrosis

**DOI:** 10.1101/2023.05.02.538835

**Authors:** Younis Hazari, Hery Urra, Valeria A. Garcia Lopez, Javier Diaz, Giovanni Tamburini, Mateus Milani, Philippe Pihan, Sylvere Durand, Fanny Aprahamia, Reese Baxter, Menghao Huang, X Charlie Dong, Helena Vihinen, Ana Batista-Gonzalez, Patricio Godoy, Alfredo Criollo, Vlad Ratziu, Fabienne Foufelle, Jan G. Hengstler, Eija Jokitalo, Beatrice Bailly-maitre, Jessica L Maiers, Lars Plate, Guido Kroemer, Claudio Hetz

**Affiliations:** Biomedical Neuroscience Institute (BNI), Faculty of Medicine, University of Chile, Santiago, Chile; FONDAP Center for Geroscience (GERO), Brain Health and Metabolism, Santiago, Chile; Program of Cellular and Molecular Biology, Institute of Biomedical Sciences, University of Chile, Santiago, Chile; Department of Biological Sciences, Vanderbilt University, Nashville, TN 37240, USA; Metabolomics and Cell Biology Platforms, UMS AMMICa Gustave Roussy Cancer Campus, Villejuif 94805, France.; Centre de Recherche des Cordeliers, Université de Paris, Sorbonne Université, INSERM U1138, Institut Universitaire de France, Paris 75006, France; Department of Medicine, Indiana University School of Medicine, Indianapolis, IN, 46202, USA; Deparment of Biochemisry and Molecular Biology, Indiana University School of Medicine, Indianapolis, IN, 46202, USA; Electron Microscopy Unit, Institute of Biotechnology, Helsinki Institute of Life Science, University of Helsinki Helsinki, Finland; Instituto de Investigación en Ciencias Odontológicas (ICOD), Facultad de Odontología, Universidad de Chile, Santiago, Chile; Leibniz Research Centre for Working Environment and Human Factors at the Technical University of Dortmund (IfADo), Ardeystrasse 67, 44139 Dortmund, Germany; Centre de Recherche des Cordeliers, INSERM, Sorbonne Université, Université de Paris, 75006 Paris, France; Institute of Cardiometabolism and Nutrition, ICAN, Assistance Publique-Hopitaux de Paris, Paris, France; Sorbonne Université, Assistance Publique-Hôpitaux De Paris, Hôpital Pitié-Salpêtrière, 75013 Paris, France; Université Côte d’Azur, INSERM, U1065, C3M, 06200 Nice, France; Department of Chemistry, Vanderbilt University, Nashville, TN 37240, USA; Institut du Cancer Paris CARPEM, Department of Biology, Hôpital Européen Georges Pompidou, AP-HP, Paris, France; Buck Institute for Research on Aging, Novato, CA, 94945, USA

**Keywords:** collagen, liver fibrosis, UPR, IRE1, PDIA1/P4HB

## Abstract

Collagen is one the most abundant proteins and the main cargo of the secretory pathway, contributing to hepatic fibrosis and cirrhosis due to excessive deposition of extracellular matrix. Here we investigated the possible contribution of the unfolded protein response, the main adaptive pathway that monitors and adjusts the protein production capacity at the endoplasmic reticulum, to collagen biogenesis and liver disease. Genetic ablation of the ER stress sensor IRE1 reduced liver damage and diminished collagen deposition in models of liver fibrosis triggered by carbon tetrachloride (CCl_4_) administration or by high fat diet. Proteomic and transcriptomic profiling identified the prolyl 4-hydroxylase (P4HB, also known as PDIA1), which is known to be critical for collagen maturation, as a major IRE1-induced gene. Cell culture studies demonstrated that IRE1 deficiency results in collagen retention at the ER and altered secretion, a phenotype rescued by P4HB overexpression. Taken together, our results collectively establish a role of the IRE1/P4HB axis in the regulation of collagen production and its significance in the pathogenesis of various disease states.

## Introduction

Collagen constitutes ∼30% of the total human proteome, it is synthesized at the endoplasmic reticulum (ER) involving a highly complex process. Deregulation in collagen biogenesis plays a critical role in various pathological conditions like osteogenesis imperfecta, fibrosis, chondrodysplasia, Ehlers–Danlos syndrome, among others [1–4]. Collagen is the principal extracellular matrix (ECM) component [5, 6], synthesized as procollagen and assembled as triple helix with (Gly-X-Y)_n_ repeats where X and Y are usually proline and hydroxyproline residues [7–10]. Proline hydroxylation, which stabilizes the triple helical conformation [11], is an enzymatic reaction catalyzed by prolyl-4-hydroxylase, a tetramer composed by two subunits of P4HA1 and two subunits of P4HB (the latter is also known as protein disulfide isomerase or PDIA1) [12, 13]. Collagen secretion also requires the ER resident collagen-specific chaperone heat shock protein 47 (Hsp47), which assists the transport of the triple helix procollagen from the ER lumen to the cis-Golgi [14, 15]. Upon liver injury, activated hepatic stellate cells (HSCs) display enhanced migration and augmented production of collagen along with other components constituting the ECM [16]. The sheer quantity of collagen production and the inherent complexity of the collagen biosynthetic pathway exert pressure on the proteostasis machinery of the secretory pathway[17, 18]. Thus, cells need to adjust their protein production capacity at multiple levels, including translation, folding, maturation, quality control, degradation, and trafficking. The ER operates as a sensing unit to monitor the efficiency of protein production, mediated by the activation of the unfolded protein response (URP), an adaptive signaling pathway that engages multiple mechanisms to cope with protein misfolding and restore proteostasis [19, 20].

The activation of the UPR is mediated by three distinct transducers, inositol requiring transmembrane kinase/ endoribonuclease 1 alpha (IRE1) and beta, PKR-like ER kinase (PERK) and activating transcription factor-6 (ATF6) alpha and beta [21]. IRE1 is a kinase and endoribonuclease that upon activation catalyzes the processing of the mRNA encoding for X-Box protein-1 (XBP1), eliminating a 26-nucleotide intron that shifts the reading frame of the mRNA [22]. This unconventional splicing event results in the expression of a potent and stable transcriptional factor known as XBP1s (for the spliced form) that translocates to the nucleus to induce the adaptive responses [22–24]. XBP1s target genes comprise factors involved in almost every aspect of the secretory pathway, including chaperones, proteins involved in ER and Golgi biogenesis, ER protein translocation, trafficking, quality control mechanisms and ER-associated degradation (ERAD) of misfolded proteins [25]. The RNase domain of IRE1 is also involved in RNA degradation (known as regulated IRE1-dependent decay or RIDD), controlling the stability of ER-localized and cytosolic mRNAs, and microRNAs [26]. IRE1 has been implicated in liver regeneration by directing hepatocyte reparative proliferation responses [27]. In addition, XBP1 expression contributes to liver regeneration after partial hepatectomy [28]. However, acute acetaminophen toxicity appears not to engage XBP1 and protects against liver toxicity mainly through activation of RIDD [29]. Interestingly, one of the few canonical RIDD substrates is the mRNA encoding collagen 6a [30], suggesting that under ER stress collagen production is reduced to alleviate ER stress generated by misfolded collagen [31]. Similarly, a few recent studies have highlighted the role of the UPR in collagen biogenesis. Activation of HSCs drives XBP1-dependent fibrogenic activity and the ectopic overexpression of XBP1s induces type I collagen expression [32]. Genome-wide transcriptomic profiling of HSCs indicated that XBP1s upregulates the collagen biosynthetic pathway, but not the transforming growth factor-beta (TGF-β) pathway [32].

Pharmacological inhibition of IRE1 with small molecules attenuates hepatic fibrosis, reduces lung fibrogenesis and limits hepatocellular carcinoma progression [33, 34]. However, the specificity of these inhibitors has been questioned, making difficult to determine the functional role of IRE1 in fibrosis [35–37]. Furthermore, blocking IRE1 activity was shown to limit the progression of simple steatosis to non-alcoholic steatohepatitis (NASH) [38]. *Xbp1* gene ablation in the liver reduces hepatic lipogenesis and leads to hypolipidemia in mice [39], contrary to other studies showing that IRE1 deletion induces hepatosteatosis and dyslipidemia due to decreased expression of microsomal triglyceride-transfer protein l [40]. Therefore, the signaling outputs controlled by IRE1 contributing to liver disease are still unclear.

Here we explored the possible contribution of IRE1 to the regulation of collagen production and secretion in an array of distinct models of liver disease including fibrosis, steatosis, high fat diet (HFD), and acute hepatotoxicity. We show that IRE1 deficiency reduces hepatocyte death in the context of acute insults and limits the deposition of collagen under persistent stress triggered by CCl_4_ or HFD. Proteomic profiling of the tissue from IRE1 depleted liver revealed that the downregulation of P4HB was a common factor in both disease models. Using cell culture models, we show that loss of IRE1 expression results in the misfolding of collagen 1 and its retention in the form of aggregates within the ER, a phenomenon reverted by P4HB overexpression. Furthermore, we found a positive correlation between *IRE1* and *P4HB* expression in human NASH samples. Taken together, our results suggest that IRE1 signaling controls collagen production in the liver possibly through regulating the expression of prolyl-4-hydroxylase and thereby defining the outcome of diseased states involving fibrosis.

## Results

### Hepatic *Ern1* <u>ablation</u> protects from CCl_4_ toxicity

To determine the contribution of IRE1 (encoded by *Ern1*) to liver disease, we generated a conditional knockout mouse using an inducible system controlled by an interferon-responsive promoter to express the Cre recombinase (MxCre) [41]. To ablate IRE1 expression in adult animals, we first used previously described floxed mice [42] to specifically delete the RNase domain, resulting in the expression of a truncated inactive protein lacking amino acids 843 to 907 (Figure 1a). Two months old mice (C57Bl6 background) were intraperitoneally injected with polyinosinic:polycytidylic acid poly-IC to induce Cre expression. Four weeks later, mice received a single dose of archetypical hepatotoxin CCl_4_ intraperitoneally at sublethal doses (1.6 g/kg) and were evaluated for liver damage after 24 hours later. As we previously reported [43], CCl_4_ treatment induced a rapid transcriptional ER stress response, as measured by the upregulation of the mRNA encoding for the ER chaperone *Bip*, the proapoptotic factor *Chop* and the ER translocation component *Sec61* (regulated by XBP1s [44]), as determined by real time PCR (Figure 1b). Importantly, this ER stress response was reduced upon knockout of IRE1 in the liver (Figure 1b). XBP1 mRNA splicing was strongly increased upon CCl_4_ administration as determined by three independent assays and was fully prevented in the liver of *Ern1^ΔR^* mice, confirming the efficiency of IRE1 deletion with the MxCre system (Figure 1c and Figure S1a). These results demonstrate that CCl_4_ triggers a rapid ER stress response in the that is mediated by IRE1.

**Figure 1.**
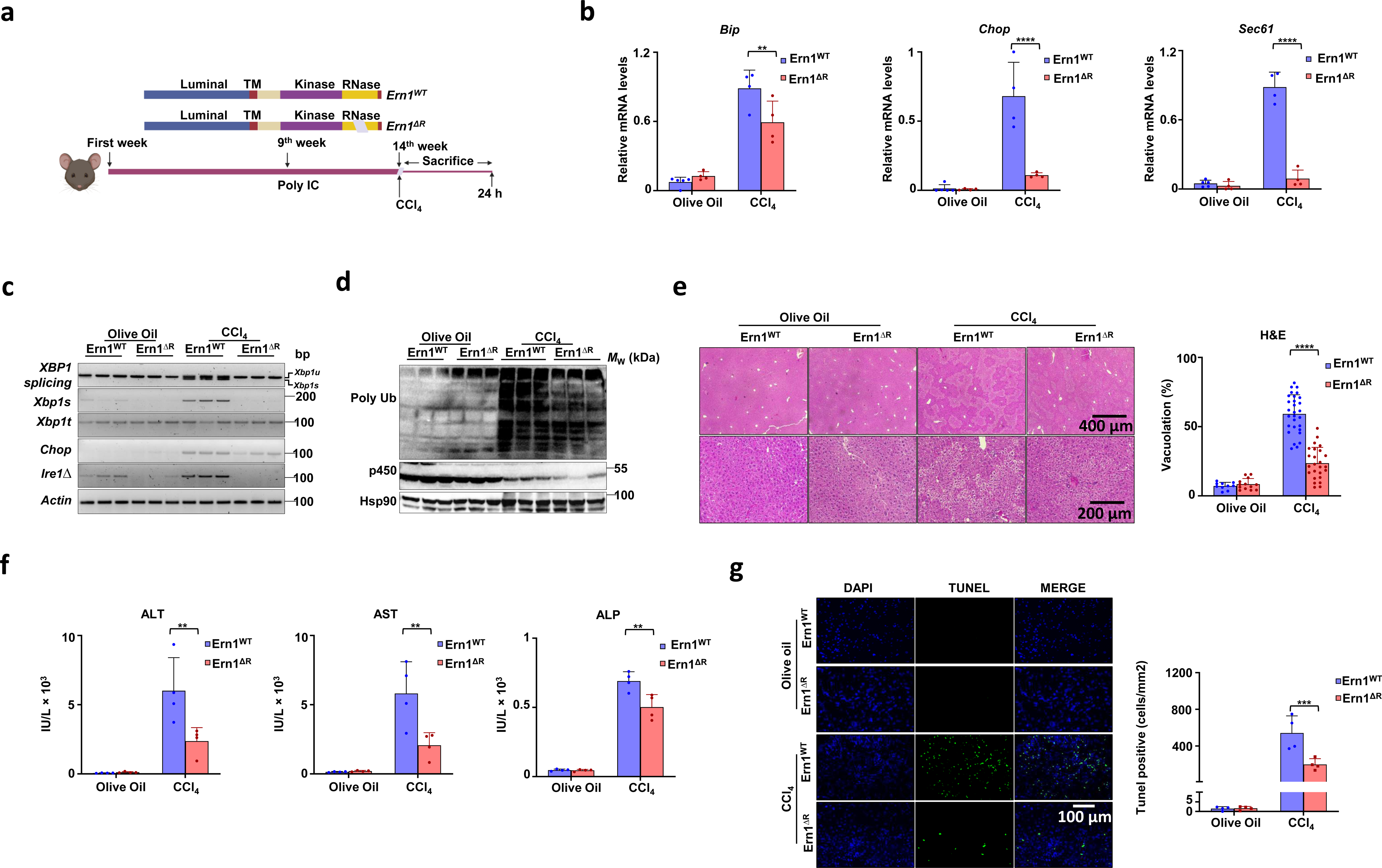
Ablation of IRE1 signaling intermits ER stress response and protects against hepatic toxicity. *Ern1* and *Ern1^ΔR^* age-matched mice littermates were treated with CCl_4_ for 24 h (n=5 for *Ern1^WT^* olive oil and n=4 per group for rest) (a) Schematic representation of IRE1showing deleted region in RNase domain and timeline of CCl_4_ treatment. (b) mRNA levels of *Bip, Chop*, and *Sec61* of *Ern1* and *Ern1^ΔR^* olive oil/CCl_4_ (1.6 g/Kg, i.p.) treated mice liver lysate after 24 h h (n=5 for *Ern1^WT^* olive oil and n=4 per group for rest). (c) Regular splicing assay for *Xbp1* mRNA in *Ern1* and *Ern1^ΔR^* olive oil/CCl_4_ (1.6 g/Kg, i.p.) 24 h treated mice liver lysate. Semi-quantitative evaluation of *Xbp1s* and *Chop* was also performed with *Xbp1t* and *Actin* as controls. To confirm the *Ern1* deletion, *Ire1Δ* measured which corresponds to the deleted mRNA region. Each well represents an individual animal (n=3 per group). (d) Immunoblotting of liver extracts using the indicated antibodies and approximate molecular weight. Representative results are shown for three individual mice of the olive oil or CCl_4_ treatment group in *Ern1* and *Ern1^ΔR^* mice after 24 h treatment. HSP90 was used as loading control. (e) H&E staining of liver sections. Representative images are shown *Ern1* and *Ern1^ΔR^* olive oil/CCl_4_ (1.6 g/Kg, i.p.) 24 h treated mice liver sections of given scale bar. Percentage loss of cytoplasm (faintly stained pink cytoplasm) or cytoplasmic vacuolation was evaluated with ImageJ software (n=4 per group, 10 to 25 images per group). (f) Serum levels of alanine aminotransferase (ALT), aspartate aminotransferase (AST) and alkaline phosphatase (ALP) after 24 h olive oil/CCl_4_ treatment *Ern1* and *Ern1^ΔR^* mice (n=4 per group). (g) *Ern1* and *Ern1^ΔR^* paraffin-embedded liver sections were stained with TUNEL (green) and DAPI (blue) (Scale bar=100 μm). Quantification of the numbers of TUNEL-positive nuclei per field in liver sections (n=4,). Statistically significant differences were determined by Two-Way ANOVA followed by Sidak’s multiple comparisons test.

We then examined general markers of proteostasis in the liver of CCl_4_ treated animals. We measured the total levels of protein ubiquitin conjugates. Consistent with reduced ER stress markers, the accumulation of polyubiquitinated proteins upon exposure of animals to CCl_4_ was attenuated in the livers of *Ern1* deleted mice (24 h treatment), suggesting improved proteostasis (Figure 1d). We also measured other components of the proteostasis network, such as autophagy, and although the pathway was induced by CCl_4_ treatment, it was not altered in *Ern1^ΔR^* mice (Figure 1Sb). In addition, analysis of oxidative stress levels using DCFDH assay indicated similar levels of reactive oxygen species (ROS) in *Ern1^WT^* and *Ern1^ΔR^* mouse liver, suggesting that CCl_4_ was equally effective in targeting this tissue (Figure S1c) [45, 46]. CCl_4_ is metabolized in the liver by cytochrome oxidase P450 enzymes, such as Cyp1a2, Cyp2e1 and Cyp3a4, generating the trichloromethyl radical CCl3*. This reaction results in a “suicide enzyme reaction” leading to the degradation of cytochrome oxidase P450 enzymes complex [47]. No significant changes were found in the P450 levels in *Ern1^ΔR^* liver samples after 24 hours of CCl_4_ administration (Figure 1d).

Next, we measured the extent of liver damage in animals exposed to CCl_4_. Hematoxylin and eosin (H&E) staining revealed extensive vacuolization and reduced cytoplasmic staining, suggesting pericentral tissue damage in *Ern1^WT^* control mice liver after 24 h after CCl_4_ administration (Figure 1e). IRE1 deficiency provided strong protection against these histopathological signs of liver damage (Figure 1e). Further, we measured markers of liver toxicity by determining the levels of alanine aminotransferase (ALT), aspartate aminotransferase (AST) and alkaline phosphatase (ALP) in blood samples [29]. Consistent with our histological analysis, IRE1 deficiency strongly reduced serum ALT, AST and ALP levels compared to *Ern1^WT^* mice (Figure 1f). Macroscopic examination revealed a more intensively speckled and coarse liver surface indicative of focal necrosis in *Ern1^WT^* mice as compared to *Ern1^ΔR^* animals (Figure 1Sd). In accordance with our previous results, analysis of liver apoptosis using the TUNEL assay indicated that *Ern1* deficiency reduces cell death upon CCl_4_ treatment (Figure 1g). Taken together, these results suggest that inhibition of IRE1 signaling protects against acute liver toxicity.

### Conditional hepatic *Ern1* deficiency limits collagen deposition

Since continuous hepatic toxicity over a prolonged time leads to enhanced secretory output with scar formation, we decided to investigate the biological consequence of *Ern1* deletion on the progression of experimental fibrosis in *Ern1^ΔR^* mouse livers. We used a protocol of chronic administration of CCl_4_ (0.4 g/kg mice 3 times per week for 12 consecutive weeks Figure 2a) Histological analysis of control mice liver showed normal lobular architecture with intact central and radiating veins based on H&E staining (Figure S2a), whereas CCl_4_ treatment resulted in disorganized liver architecture and collagen fiber deposition as measured by Masson trichrome staining (Figure 2b). In sharp contrast, *Ern1^ΔR^* mice livers showed a near 50% reduction in the amounts of collagen deposition (Figure 2b). Similar results were obtained using Sirius red staining (Figure 2b). We also observed significant reduction in ALT and AST serum levels after chronic administration of CCl_4_ to *Ern1^ΔR^* mice compared to litter mate controls (Figure 2c). The increase in the liver-to–body weight ratio observed after CCl_4_ administration was attenuated upon ablation of IRE1 expression (Figure 2d). We further confirmed a reduction in collagen I deposition in the liver of *Ern1^ΔR^* mice treated with CCl_4_ as determined with immunofluorescence staining (Figure 2e). Similar results were obtained when the amount of collagen deposition was determined colorimetrically by measuring the hydroxyproline content (Figure 2f), a key collagen post-translational modification in fibrotic livers.

**Figure 2.**
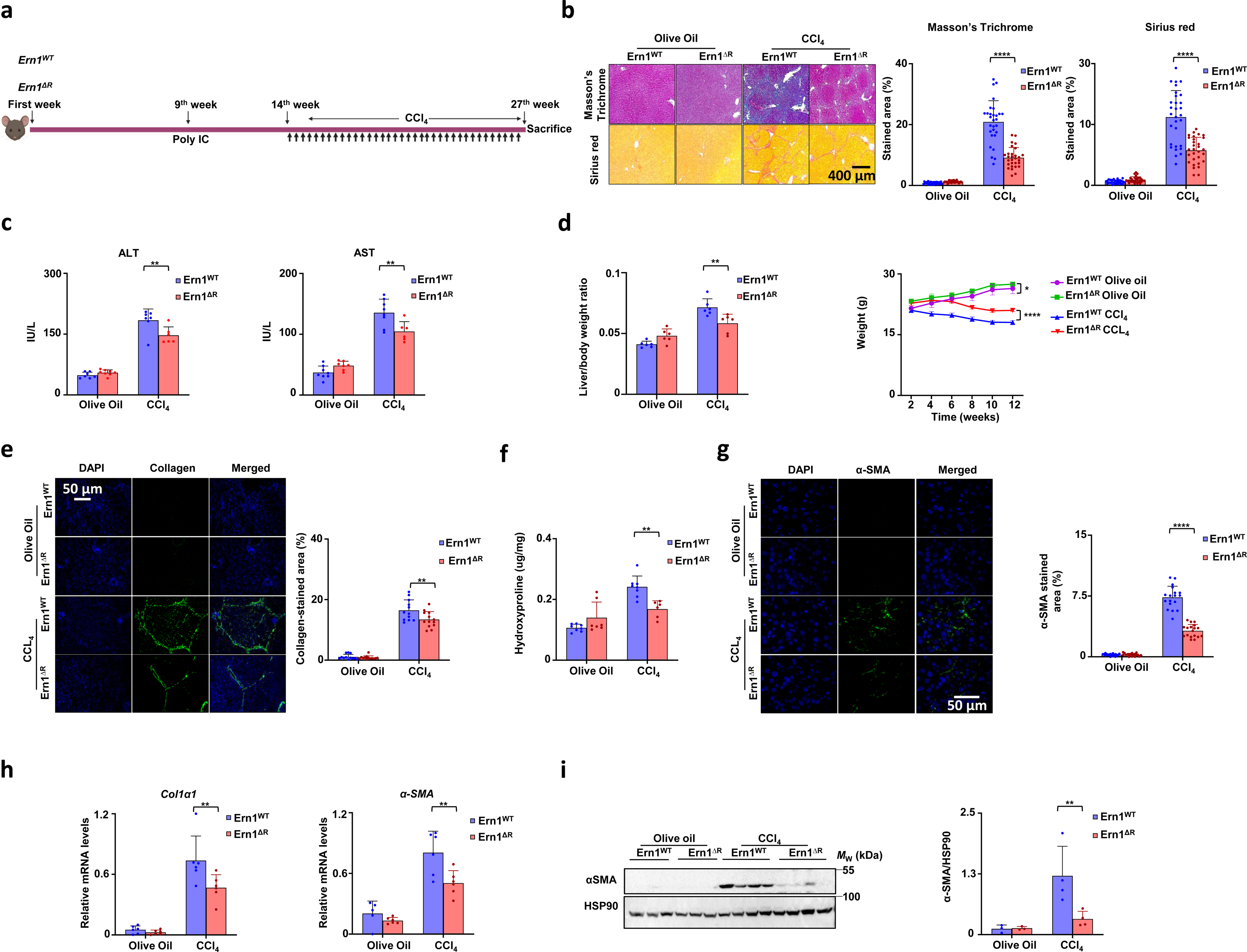
IRE1 deficiency improves chronic fibrosis with reduced collagen deposition. *Ern1* and *Ern1^ΔR^*age-matched mice littermates were treated with CCl_4_ for 3 doses per week on alternative days of CCl_4_ (0.4 g/Kg) for 12 consecutive weeks (n=7 to 8 per group) (a) Schematic representation of IRE1showing deleted region in RNase domain and timeline of CCl_4_ treatment. (b) Masson’s Trichrome and Sirius red staining of liver sections after CCl_4_ for 12 weeks (n=8 to 7 per group), scale bar=400 μm and their respective quantification of stained liver fibrosis area (percent of total area) (c) Serum levels of alanine aminotransferase (ALT) and aspartate aminotransferase (AST). (d) Liver weight was determined relative to body weight (n=6 to 7) and weekly body weight is shown. Statistical analysis was performed using two-way ANOVA with Tukey’s multiple comparison test. Mean ±S.E. is shown (n=6 to 7). (e) Immunofluorescence of CCl_4_ treated *Ern1* and *Ern1^ΔR^* mice. Liver sections were stained with collagen (green) and Dapi (blue) (n=6 to 7), scale bar=50 μm and scatter plots represent collagen-stained area measured by ImageJ. (f) Liver hydroxyproline content was determined to estimate collagen content and expressed as µg hydroxyproline per mg liver tissue (wet weight). Statistical analysis was performed using two-way ANOVA with Tukey’s multiple comparison test. Mean ±S.E. is shown (n=6 to 7). (f) Representative confocal images α-SMA (green) immunofluorescence in the liver of CCl_4_ treated *Ern1* and *Ern1^ΔR^* mice liver sections. Nuclei are stained blue. Scale bar =50 μm. Scatter plots represent percentage α-SMA positive area (n=6 to 7). (g) mRNA levels of *Col1α1* and *α-SMA* of *Ern1* and *Ern1^ΔR^* of CCl_4_ (0.4 g/Kg) for 12 consecutive weeks treated mice liver lysate. (h) Immunoblotting of liver extracts against α-SMA antibody with approximate molecular weight. Representative results are shown for three individual mice of the olive oil or 4 individual mice of CCl_4_ treatment group in *Ern1* and *Ern1^ΔR^* mice. HSP90 was used as loading control. Statistically significant differences were determined by Two-Way ANOVA followed by Sidak’s multiple comparisons test.

We next measured the levels of other factors involved in the progression of liver fibrosis. The CCl_4_-induced expression of α-SMA (Acta2), a classic marker for HSC activation was reduced in *Ern1^ΔR^* mice liver samples, as determined by immunofluorescence microscopy (Figure 2g). We also found reduced upregulation of the mRNA levels encoding for *α-SMA* and *Col1a1* in *Ern1^ΔR^* livers (Figure 2h). To complement these findings, we analyzed α-SMA levels in fibrotic livers by immunoblotting, confirming reduced upregulation upon CCl_4_ treatment in *Ern1^ΔR^* mice (Figure 2i). The expression of transforming growth factor-β (*TGF-β*), a major fibrogenic cytokine, also showed reduced levels in the liver of CCl_4_ treated *Ern1^ΔR^* animals (Figure S2b). Furthermore, matrix remodeling genes including tissue inhibitor matrix metalloproteinase 1 (*Timp1*) and matrix metalloproteinase-2 (*MMP2*) showed significant reduction in CCl_4_ treated *Ern1^ΔR^* mice, suggesting that the pathways involved in fibrosis are attenuated upon targeting IRE1 (Figure S2b). Thus, IRE1 deficiency in liver leads to persistent altered fibrotic cascade with overall reduction in collagen deposition.

### Hepatic *Ern1* deletion downregulates P4hb *in vivo*

IRE1 regulates proteostasis through the activation of the UPR transcription factor XBP1, in addition to controlling the stability of certain mRNAs through RIDD. However, IRE1-regulated changes in the transcriptome largely depend on experimental perturbators and cell types [26]. To determine possible molecular mechanisms underlying the protective effects of IRE1 deficiency, we performed mass spectrometry-based quantitative proteomics of liver samples from *Ern1^WT^* and *Ern1^ΔR^* animals treated or not with CCl_4_ (animals from figure 1). Intriguingly, proteomic assessment at basal levels revealed the downregulation of 10 proteins due to IRE1 depletion at basal levels, highlighting P4HB (Figure 3a and 3b), a limiting factor in collagen maturation. Furthermore, gene set enrichment revealed altered levels of collagen transportation factors involved in ER exit, among other proteostasis processes related to the secretory pathway (Figure 3c and 3d). Analysis of liver samples from CCl_4_-treated animals confirmed a consistent downregulation of P4HB in IRE1 deficient livers (Figure S3a and S3b). IRE1 deficiency in CCl_4_ treated animals also altered components of the secretory pathway, in addition to metabolic processes and detoxification pathways (Figure S3c and S3d). Taken together, these results suggest P4HB dependency on the IRE1 to regulate the proline hydroxylation of collagen (Figure S3c and S3d).

**Figure 3.**
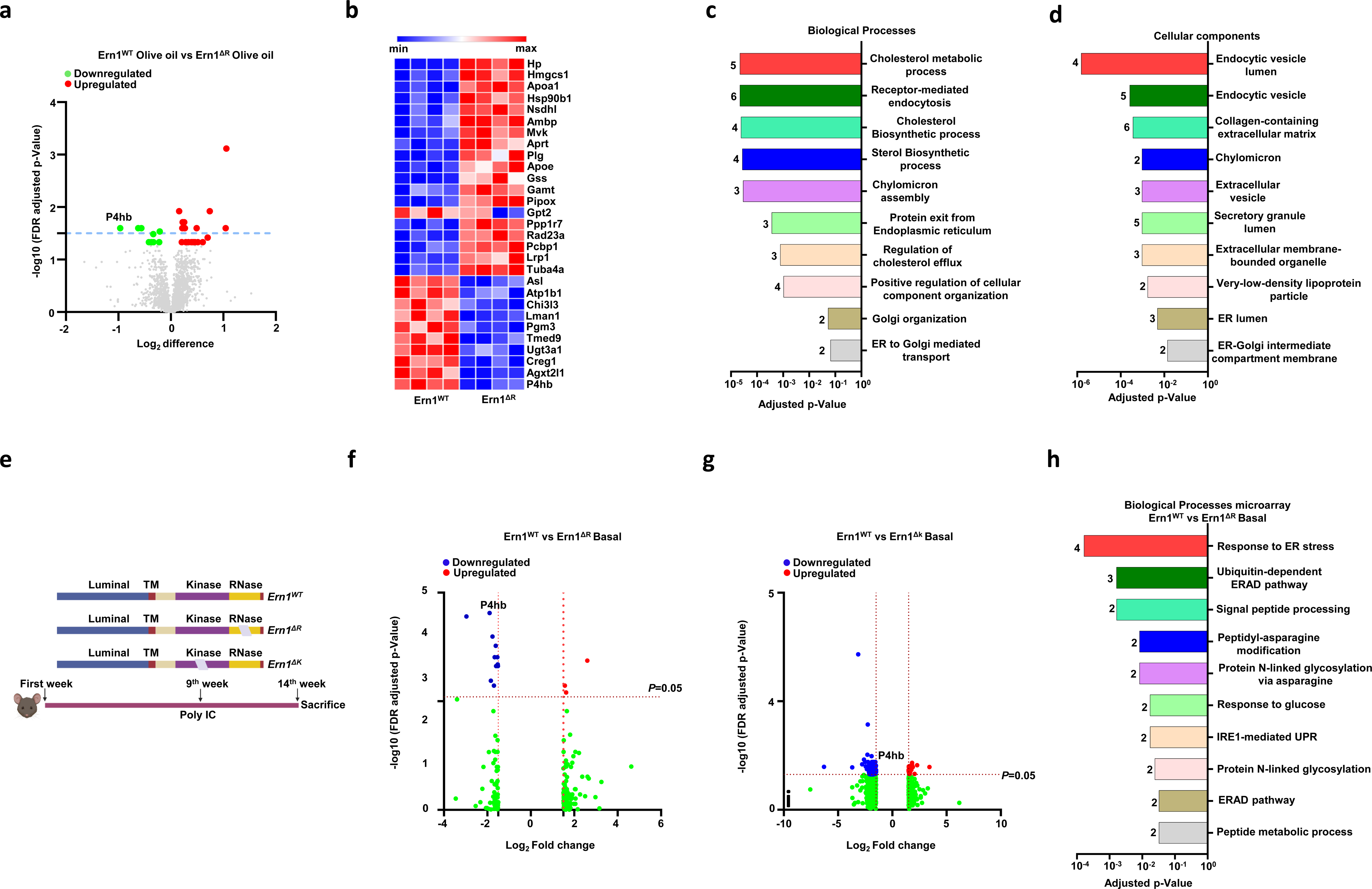
Hepatic *Ern1* deletion regulates protein expression in the liver. (a) Volcano plot of proteomic analysis of olive oil treated *Ern1* and *Ern1^ΔR^* mice liver. The x-axis represents the logarithmic fold change levels in the *Ern1^ΔR^* mice liver relative to the *Ern1* mice liver and y-axis represents the negative decade logarithm of FDR adjusted P-values (n=4). Statistical analysis of unpaired t test with 5% False discovery rate using Two-stage step-up Benjamini. Krieger and Yekutiele method assuming individual variance for each row. Selected hits shown have q value ≤ 0.05 and P-value ≤ 0.05. Green dots indicate the downregulated proteins and red dots indicate the upregulation of proteins. (b) The heatmap represents the 10 downregulated (blue) and 19 upregulated proteins (red) from the *Ern1* and *Ern1^ΔR^*mice liver. Each lane represents individual mice and color intensity denotes the abundance of the protein. (c) Functional classification of proteomic hits according to the Gene Ontology (GO) annotation. Graph shows significantly enriched GO terms of biological processes. The number of Genes associated with each biological process is indicated by the respective bar. in the parenthesis. (d) Functional classification of proteomic hits according to the Gene Ontology (GO) annotation. Graph shows significantly enriched GO terms of cellular components. The number of Genes associated with each biological process is indicated by the respective bar. (e) Schematic representation of IRE1showing deleted region in RNase and Kinase domain with timeline of experiment. (f) Volcano plot of microarray analysis of olive oil treated *Ern1* and *Ern1^ΔR^* mice liver. The x-axis represents the logarithmic fold change levels in the *Ern1^ΔR^* mice liver relative to the *Ern1* mice liver and y-axis represents the negative decade logarithm of FDR adjusted P-values. Statistical analysis of unpaired t test with 5% False discovery rate using Two-stage step-up Benjamini. Krieger and Yekutiele method assuming individual variance for each row. Selected hits shown have q value ≤ 0.05 and P-value ≤ 0.05 (n=3). Blue dots indicate the downregulated proteins and red dots indicate the upregulation of proteins. (g) Volcano plot of microarray analysis of olive oil treated *Ern1* and *Ern1^Δk^* mice liver. The x-axis represents the logarithmic fold change levels in the *Ern1^Δk^* mice liver relative to the *Ern1* mice liver and y-axis represents the negative decade logarithm of FDR adjusted P-values. Statistical analysis of unpaired t test with 5% False discovery rate using Two-stage step-up Benjamini. Krieger and Yekutiele method assuming individual variance for each row. Selected hits shown have q value ≤ 0.05 and P-value ≤ 0.05 (n=3). Blue dots indicate the downregulated proteins and red dots indicate the upregulation of proteins. (h) Functional classification of microarray analysis hit in the *Ern1* and *Ern1^ΔR^* mice liver according to the Gene Ontology (GO) annotation. Graph shows significantly enriched GO terms of biological processes. The number of Genes associated with each biological process is indicated by the respective bar.

We next performed a global gene expression analysis using microarray and compared relative mRNA levels in *Ern1^ΔR^* and control livers. The rationale behind this evaluation was to determine the effects of IRE1 deficiency in the liver at the unstressed state, and then to validate findings we used a second model for IRE1 deficiency instead of the RNase domain (*Ern1^ΔR^*) the kinase domain is ablated (*Ern1^ΔK^*) (Figure 3e) [48]. Remarkably, comparison of *Ern1^ΔR^* and *Ern1^WT^* mouse liver samples indicated the downregulation of P4HB mRNA in IRE1 deficient animals (Figure 3f and 3g). Functional enrichment analysis of *Ern1^ΔR^* microarray data revealed alterations in posttranslational modifications pathways, ER protein processing and maturation and other proteostasis processes (Figure 3h). Overall, the only common hit between the proteomic and microarray analysis was P4HB, an observation that guided our further validation experiments (Figure S3e and S3f).

Based on the results obtained in the proteomic analysis, we measured P4HB levels using immunoblotting. We found reduced levels of P4HB under resting conditions in *Ern1^ΔR^ and Ern1^ΔK^* mice with respect to IRE1-sufficient controls (Figure 4a and 4b). Real time PCR analysis corroborated reduced *P4hb* mRNA levels in both models of IRE1 deficiency (Figure 4c and 4d). Similar observations were upon chronic exposure of *Ern1^ΔR^* mice to CCl_4_ (Figure 4e), validating our proteomic results. Overall, these results indicate that IRE1 deficiency in the liver results on the downregulation of P4HB at the mRNA and protein level.

**Figure 4.**
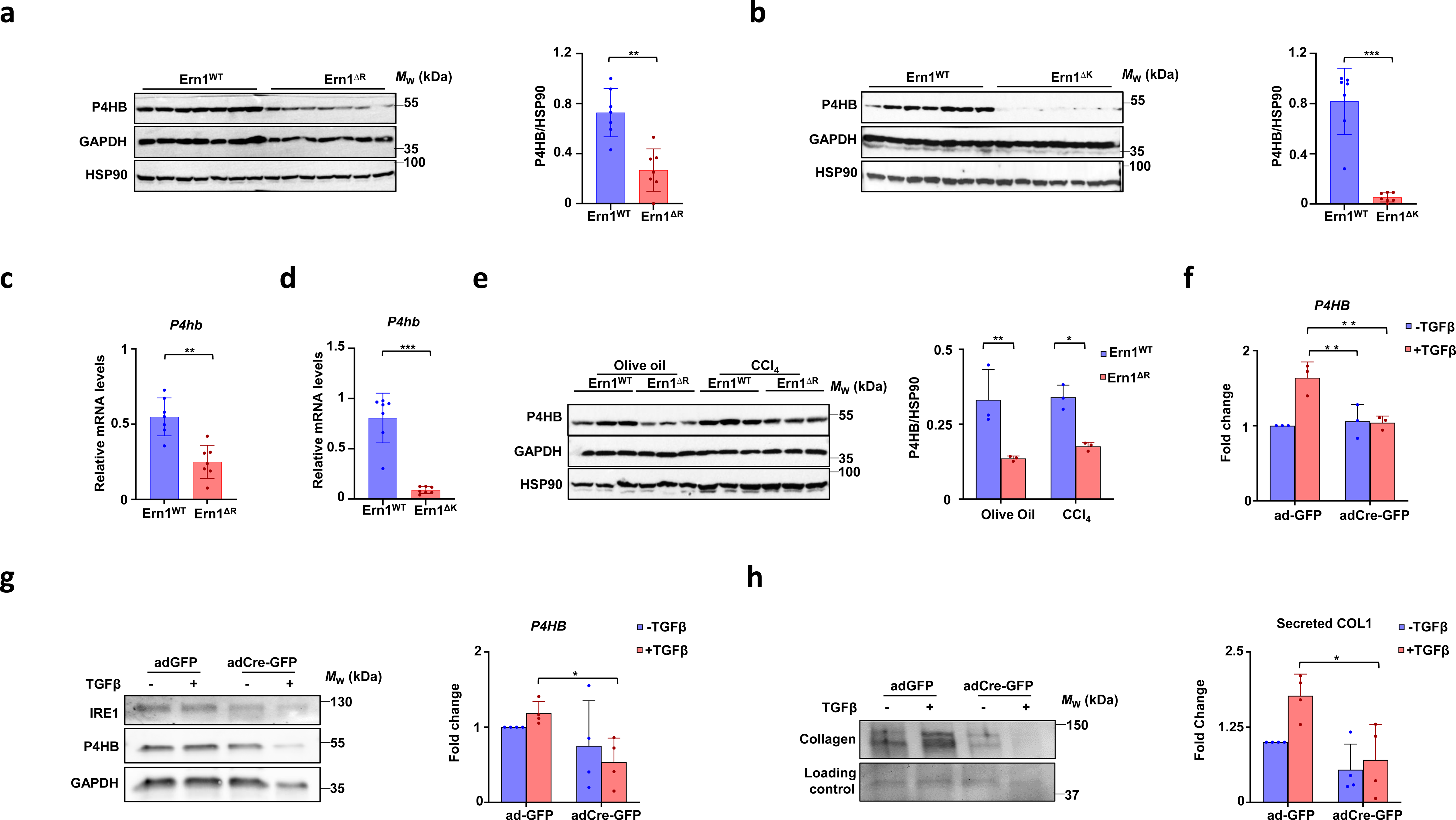
IRE1 deficiency in livers and HSCs downregulates P4HB levels. (a) Immunoblotting of liver extracts for P4HB antibody with approximate molecular weight. Representative results are shown for 7 individual mice of *Ern1* and *Ern1^ΔR^* mice liver. HSP90 was used as loading control. (b) Immunoblotting of liver extracts for P4HB antibody with approximate molecular weight. Representative results are shown for 7 individual mice of *Ern1* and *Ern1^ΔK^* mice liver. HSP90 was used as loading control. (c) mRNA level of *P4hb* of *Ern1* and *Ern1^ΔR^* mice liver lysate. *Actin* was used for normalization. P value was calculated using Mann-Whitney test. (d) mRNA level of *P4hb* of *Ern1* and *Ern1^ΔK^* mice liver lysate. *Actin* was used for normalization. The P value was calculated using Mann-Whitney test. (e) Immunoblotting of liver extracts using the indicated antibodies with approximate molecular weight. Representative results are shown for three individual mice of the olive oil or CCl_4_ treatment group in *Ern1* and *Ern1^ΔR^* mice. GAPDH was used as loading control. Statistical analysis was performed using Mann-Whitney test. (f) mRNA levels of *P4hb of Ern1* and *Ern1* floxed mice of isolated and activated HSCs after 2 ng/mL of TGF-β for 24 hours. P value was calculated by one-way ANOVA followed by Tukey’s multiple comparisons (N=3). (g) Immunoblotting of activated HSC lysate isolated from *Ern1* and *Ern1* floxed mice probed for the IRE1 and P4HB shown against approximate molecular weight. GAPDH was used as loading control. P value was calculated by one-way ANOVA followed by Tukey’s multiple comparisons (N=4). (h) Immunoblotting of media collected from activated HSC lysate isolated from *Ern1* and *Ern1* floxed mice probed for secreted collagen shown against approximate molecular weight. Total protein staining of the conditioned media was used as loading control. P value was calculated by one-way ANOVA followed by Tukey’s multiple comparisons (N=4).

Since HSC are the main collagen synthesizing cells, we decided to isolate primary HSCs from the IRE1 floxed mice and subsequently activated them with 2 ng/mL of TGF-β for 24 hours. To ablate IRE1, we delivered adenoviruses to express Cre *in vitro* (adCre-GFP). TGF-β driven HSC activation resulted in an increase in P4HB mRNA levels, whereas IRE1 ablation prevented this upregulation (Figure 4f). Consistently, protein levels of P4HB were also reduced (Figure 4g). Remarkably the augmented secretion of collagen into the culture supernatants induced by TGF-β was prevented upon targeting of IRE1 in primary HSCs (Figure 4h). Altogether, our results indicate that IRE1 deficiency in the liver results in the downregulation of P4HB, correlating with reduced collagen secretion.

### Hepatic *Ern1* deficiency suppresses the progression of liver steatosis

Our results suggest that IRE1 signaling contributes to liver damage and fibrosis. Driven by these findings, we evaluated the possible involvement of the IRE1 in steatohepatitis as chronic liver disease due to NASH is considered to be main cause of liver transplantation [49–51]. For this purpose, we exposed *Ern1* and *Ern1^ΔR^* mice to high fat diet (HFD) for 12 consecutive weeks to induce the experimental steatosis (Figure 5a). IRE1 deficiency resulted in significant decrease in liver weight and visceral fat when compared with WT mice fed with HFD (Figure 5b). The serum levels of ALT, AST and hepatic TG were also attenuated in *Ern1^ΔR^* animals, suggesting reduced liver damage in the HFD model (Figure 5c). Liver histology of *Ern1^ΔR^* HFD fed mice revealed intact liver architecture and lower lipid accumulation compared to wild-type mice, indicating reduced steatosis (Figure 5d). In addition, the deposition of collagen was also attenuated in *Ern1^ΔR^* HFD fed mice compared to *Ern1^WT^*controls (Figure 5d).

**Figure 5.**
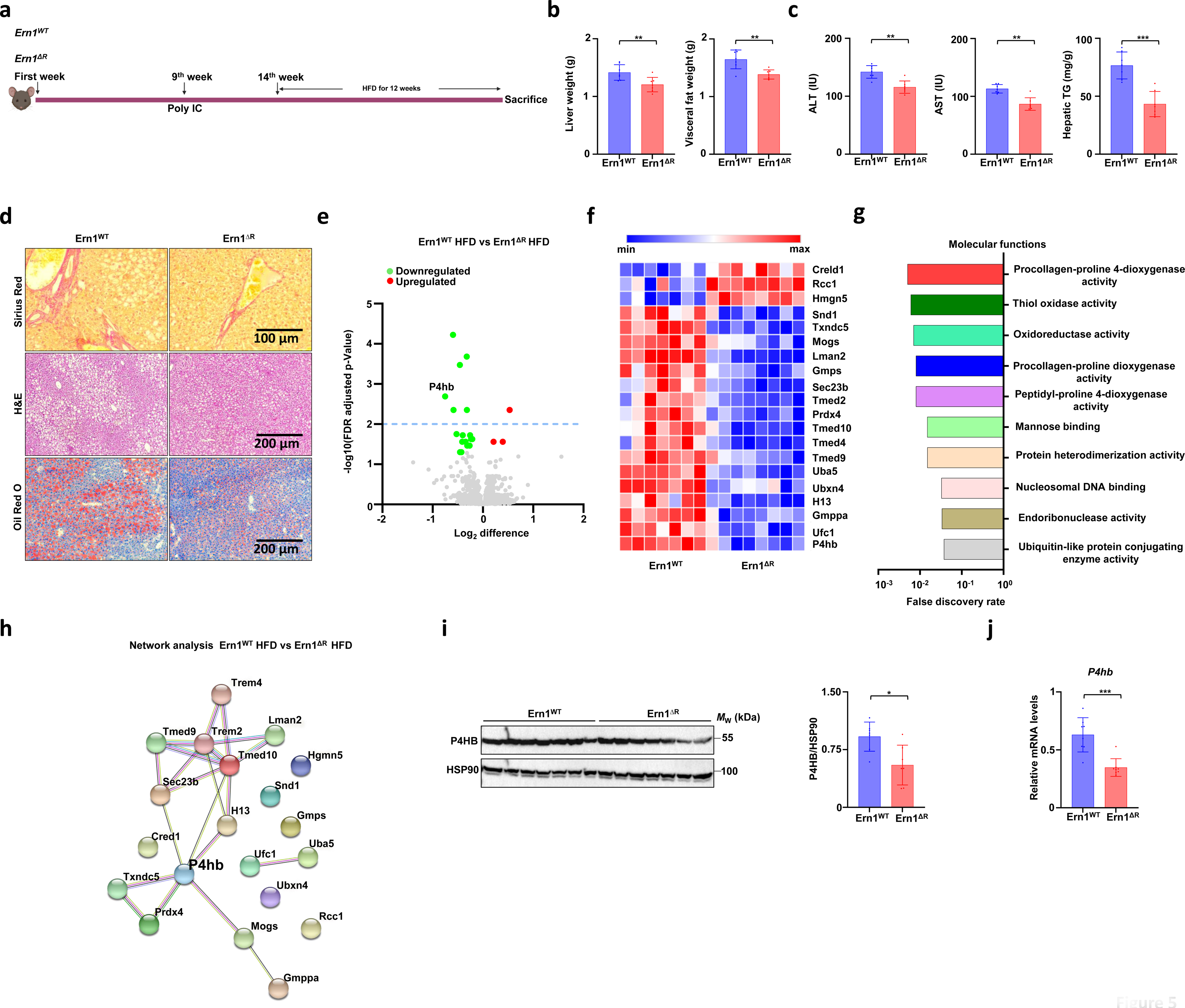
IRE1 deficiency delimits steatohepatitis in mice. (a) Schematic representation of IRE1showing deleted region in RNase domain and timeline of HFD mice model. (b) Comparative liver weight and visceral fat content of *Ern1* and *Ern1^ΔR^*mice liver fed with HFD for 12 consecutive weeks (n=7). (c) Serum levels of ALT and AST, and hepatic triglyceride content (n=7). (d) Representative photomicrograph of Sirius red staining, H&E staining, and oil red staining of HFD fed *Ern1* and *Ern1^ΔR^*mice liver for 12 consecutive weeks with denoted scale bar. (e) Volcano plot of proteomic analysis of *Ern1* and *Ern1^ΔR^* mice liver fed with HFD for 12 consecutive weeks. The x-axis represents the logarithmic fold change levels in the *Ern1^ΔR^* mice liver relative to the *Ern1* mice liver fed with HFD for 12 consecutive weeks and y-axis represents the negative decade logarithm of FDR adjusted P-values (n=7). Statistical analysis of unpaired t test with 5% False discovery rate using Two-stage step-up Benjamini. Krieger and Yekutiele method assuming individual variance for each row. Selected hits shown have q value ≤ 0.05 and P-value ≤ 0.05. Green dots indicate the downregulated proteins and red dots indicates the upregulation of proteins (n=7 for *Ern1 (*one sample was not included because the sample in this channel showed lower abundance for unnormalized peptide) and n=8 for *Ern1^ΔR^*). Green dots indicate the downregulated proteins and red dots indicate the upregulation of proteins. (f) The heatmap represents the 29 downregulated proteins and 2 upregulated proteins from the *Ern1* and *Ern1^ΔR^*mice liver. Each lane represents individual mice and color intensity denotes the abundance of the protein. (g) Functional classification of proteomic hits according to the Gene Ontology (GO) annotation. Graph shows significantly enriched GO terms of biological processes. The number of Genes associated with each biological process is indicated in the parenthesis. (h) Network analysis of P4HB interaction with identified genes related to ER to Golgi transport and collagen synthesis and secretion. The nodes represent the proteins and the edges interactions. The thickness of the edges indicates the extent of interactions, which does not necessarily mean physical binding. (i) Immunoblotting of liver extracts for P4HB antibody with approximate molecular weight. Representative results are shown for 7 individual mice of *Ern1* and *Ern1^ΔR^* mice liver fed with HFD for 12 consecutive weeks. HSP90 was used as loading control. (j) mRNA level of *P4hb* of *Ern1* and *Ern1^ΔR^* mice liver lysate fed with HFD for 12 consecutive weeks. *Actin* was used for normalization. Statistically significant differences were determined by Mann-Whitney test.

We then performed proteomic analysis of the liver of *Ern1^ΔR^* animals undergoing HFD. We found 31 hits to be significantly altered in the liver of HFD fed mice. Again, P4HB was significantly downregulated in *Ern1^ΔR^* animals (Figure 5e and 5f). Gene set enrichment analysis also revealed down-regulation of 4-hydroxyproline metabolic process with reduced expression of PRDX4 (Figure 5g and 5h), other important protein involved in the hydroxylation of prolyl residues in pre-procollagen and its metabolism [52]. Analysis of P4HB expression levels in the liver of animals exposed to HFD confirmed the downregulation of P4HB at mRNA and protein levels (Figure 5i and 5j). Overall, these results suggest that IRE1 deficiency protects the liver against pathological conditions, correlating with the downregulation of P4HB expression, a key factor involved in collagen maturation and protein folding at the ER.

Finally, we also performed a global metabolomic profiling of the livers from IRE1 deficient mice. In animals exposed to HFD, heat map and pathway analysis indicated alterations in the ascorbate and glycine metabolism (Figure S4a, S4b and S4c), which are both important for collagen synthesis. The key precursor of ascorbate, glucuronate accumulated in the *Ern1^ΔR^* liver samples (Figure S4d). We also measured metabolite levels in the liver of CCl_4_-treated animals. Heatmap and pathway analysis revealed a prominent alteration in the glutamate and proline metabolism pathways (Figure S4e, S4f and S4g). *Ern1^ΔR^* CCl_4_ treated liver samples showed clear accumulation of key metabolites like ornithine and 4-hydroxyproline (Figure S4h), indicative of possible reduction in the collagen synthesis [53]. The overall alteration of the different metabolites in HFD and CCl_4_ mice models indicate that IRE1 ablation in the liver affects metabolic pathways involved in collagen synthesis (Figure S4i).

### IRE1 regulates collagen production in vitro

Having established that IRE1 ablation reduces the P4HB levels and collagen secretion in primary HSCs, we generated *ERN1* knockout (KO) cells using CRISPR-Cas9 in U20S cells, a well-studied collagen synthesizing cells (Figure 6a) [54, 55]. The total levels of P4HB were also reduced at the protein levels in IRE1 deficient cells (Figure 6a). In addition, we found a significant downregulation of the P4HB mRNA levels upon targeting IRE1 expression (Figure 6b). A similar decrease in the P4HB levels were observed when IRE1 was ablated in the hepatocyte cell lines Hepa1-6 cells and Huh-7 (Figure 6a and 6b).

**Figure 6.**
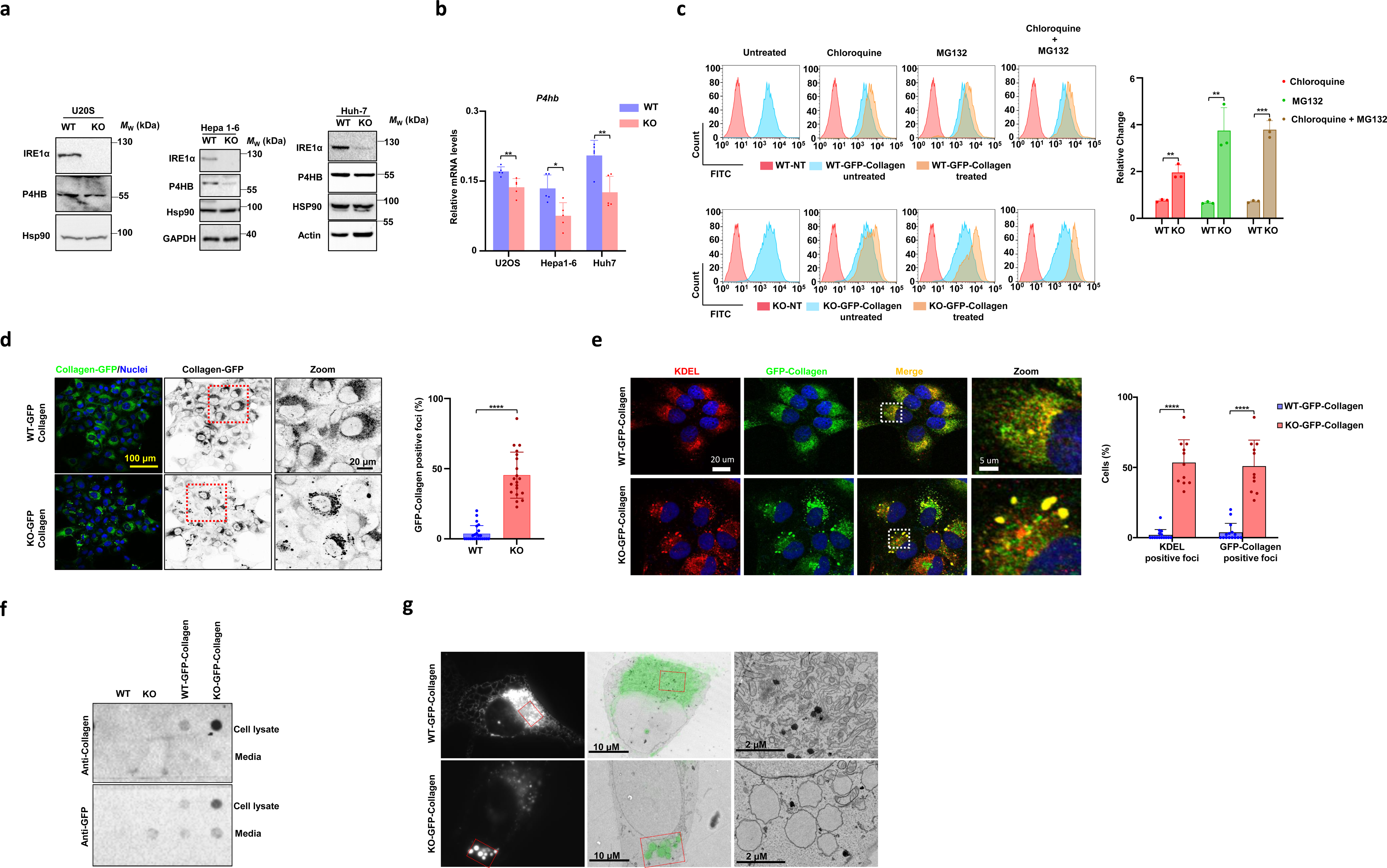
ER retention of collagen in absence of IRE1. (a) Real time qPCR of P4HB mRNA levels in U20S (single clone) (n=5), Hepa1-6 (single clone) (n=5), and Huh7 (pool) (n=5) IRE1null CRISPR-Cas9 (KO, red) and their corresponding CRISPR-Cas9 control (WT, blue). using two-tailed unpaired Student’s *t*-tests. Mean ± SEM are shown. *P < 0.05; **P < 0.01; ****P < 0.0001.(b) Immunoblots of P4HB with indicative loading controls in U20S Hepa1-6 and Huh7 IRE1null CRISPR-Cas9 (KO) and their corresponding CRISPR-Cas9 control (WT). (c) Representative histograms of GFP-collagen fluorescence in the presence or absence of chloroquine (autophagy inhibitor, 50 μm) and MG132 (proteasome inhibitor, 50 μm) for 16 h. Quantification of relative change in the median fluorescence intensity with respect to the untreated. Results are mean ± SEM of 3 independent experiments. Statistical analysis was performed using two-tailed unpaired Student’s *t*-tests. Mean ± SEM are shown (d) GFP-collagen cell’s nucleus was stained with the Dapi and GFP-collagen punta’s were quantified using ImageJ software of 3 independent experiments (n=574 CCT-collagen cells and n=639 KO-collagen cells). Statistical analysis was performed using two-tailed unpaired Student’s *t*-tests. Mean ± SEM are shown. (e) GFP-collagen cell’s nucleus was stained with the Dapi, ER with KDEL and GFP-collagen punta’s were quantified using ImageJ software of 2 independent experiments (n=372 CCT-collagen cells and n=363 KO-collagen cells). Statistical analysis was performed using two-tailed unpaired Student’s t-tests. Mean ± SEM are shown. (f) Filter-trap assay for collagen and GFP antibody. (g) Represents the CLEM of the WT-GFP-collagen and KO-GFP-collagen with given scale bar.

To determine possible molecular consequences of IRE1 deficiency to collagen biogenesis and the functional relationship with P4HB, we then generated a stable U2OS cell line expressing GFP-tagged human collagen 1A1 (COL1A1) by sorting cells with similar average GFP levels (Figure S5a) [48]. IRE1 deficiency was confirmed by measuring *xbp1* mRNA splicing in cells treated with tunicamycin (Figure S5b). Ablation of IRE1 expression increased the levels of intracellular collagen-GFP as compared with control cells (Figure 6c). Because misfolded collagen is degraded by the autophagy and the proteasomes [56], we measured the expression of GFP-collagen by FACS after treating cells with MG132 (which inhibits the proteasome) or chloroquine (which blocks final step of autophagy). Treating with MG132 profoundly increased the GFP fluorescence in the IRE1 deficient cells, whereas chloroquine showed very mild effects (Figure 6c).

Confocal microscopy analysis confirmed the presence of significantly higher GFP-collagen puncta in IRE1 KO cells (Figure 6d). Most of the GFP-collagen signal in these cells colocalized with the ER marker KDEL, as measured by immunofluorescence (Figure 6e), indicative of intracellular accumulation of immature collagen in early secretory pathway compartments. Based on these findings, we then reasoned that GFP-collagen forms intracellular aggregates more profoundly in IRE1 KO cells compared with the corresponding WT controls. Thus, we performed a filter-trap assay and probed for both collagen and GFP. We found increased amounts of high-molecular weight collagen-containing complexes in IRE1 KO cell lysates than in controls (Figure 6f). One of the adaptive responses regulated by IRE1/XBP1 under ER stress is the induction of ER and Golgi biogenesis. Consistent with this, staining with ER-tracker BODIPY followed by FACS analysis indicated a significant decrease in the staining in IRE1 KO cells overexpressing collagen when compared with CRISPR-controls (Figure S5c). Consistent with these results, visualization of cells with transmission electron microscopy (TEM) indicated decreased ER content and evident ER dilation (a sign of ER stress) (Figure S5d). Using correlative light electron microscopy (CLEM) we were able to localize the GFP-collagen within swallow and rounded ER, a phenotype seen more frequently in IRE1 KO (Figure 6g). Notably, many of the large GFP-collagen positive structures were found accompanied with budding profiles and vehicles suggesting that they are arrested to the ER exit sites. Taken together, these results indicate that ablation of IRE1 expression results in the misfolding and intracellular accumulation of collagen inside the cell, reducing its secretion.

### P4HB overexpression improves collagen biogenesis in IRE1 deficient cells

Ascorbate is essential for the enzymatic activity of prolyl hydroxylase to hydroxylate the prolyl residue modified in procollagen before triple helix formation. Stimulation of prolyl hydroxylation due to addition of ascorbate is known to increase collagen secretion [57–60]. We found that the reduction in GFP-collagen expression in IRE1 KO cells was more profound in cells treated with 5 μg/mL ascorbate when compared with GFP-collagen expressing CRISPR-control cells and remained constant with higher concentrations of ascorbate (Figure S6a).

Next, we tested whether overexpression of P4HB would rescue the collagen defects observed in IRE1 deficient cells. P4HB (PDIA1) is a member of a family of ER resident protein disulfide isomerases (PDIs), which include PDIA3 (ERp57) and PDI4 (ERp72) among others, essential factors for ER proteostasis and folding [61, 62]. We transiently expressed P4HB, ERp57 or Erp72 in IRE1 KO cells in the presence or absence of ascorbate. Overexpression of P4HB, ERp57 or ERp72 increased GFP-collagen signal, whereas on addition of ascorbate, GFP-collagen in P4HB overexpressing cells considerably diminished the accumulation of intracellular GFP-collagen (Figure 7a). The changes observed due to ERp57 and ERp72 on ascorbate addition were not significant (Figure 7a). We then measured the effects of overexpressing P4HB using immunoblotting analysis in presence or absence of ascorbate. Remarkably, P4HB overexpression reduced the accumulation of intracellular collagen in IRE1 KO cells, but also in cells treated with ascorbate (Figure 7b). We also evaluated the protein levels of several ER resident foldases and cofactors, including calnexin, calreticulin and ERO1A which remained unaltered in IRE1 KO cells (Figure S6b). We then performed filter-trap assay which revealed a considerable decrease in the GFP-collagen aggregates when P4HB was expressed in IRE1 KO cells (Figure 7c). In addition, analysis of intracellular GFP-collagen puncta (determined by confocal microscopy) indicated reduced accumulation in cells expressing V5-tagged P4HB (Figure 7d and S6c). Based on these results, we performed immunoprecipitation experiments in U2OS cells expressing GFP-collagen. We observed a basal interaction between endogenous P4HB and GFP-collagen (Figure S6d). Interestingly, when we performed similar experiments in IRE1 KO cells, a more pronounced interaction between P4HB and GFP-collagen was observed, suggesting higher levels of misfolded collagen (Figure S6d). Indeed, the electrophoretic migration of GFP-collagen was altered in IRE1 KO cells, suggesting altered maturation process. Taken together, these results indicate that the sole expression of P4HB in IRE1 null cells reduced the abnormal intracellular deposition of collagen implying that this PDI family member is one of the major effectors of IRE1 signaling in cells expressing collagen.

**Figure 7.**
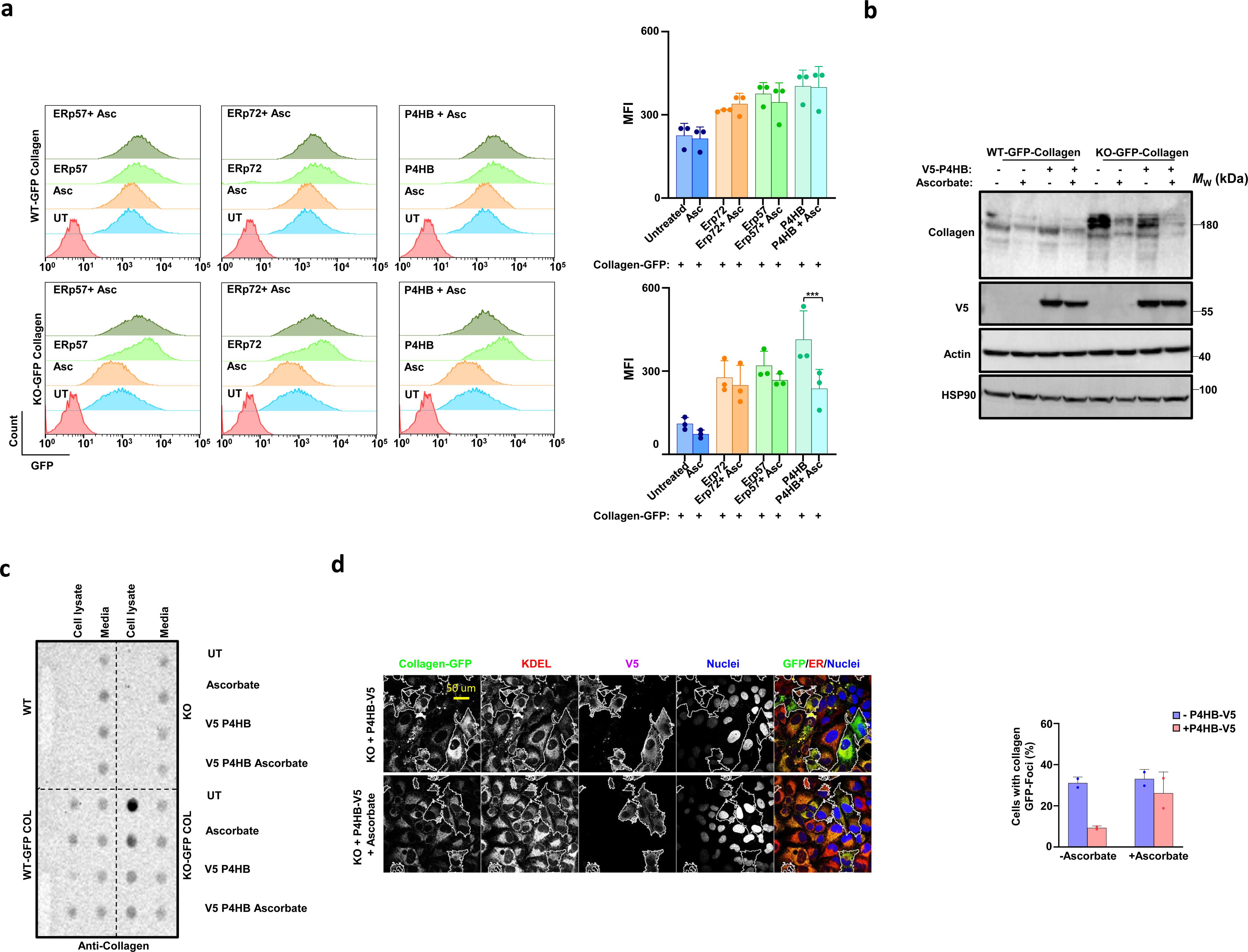
P4HB overexpression rescues collagen biogenesis due to IRE1 deficiency in cell culture. (a) Representative half offset histograms of GFP-collagen fluorescence post 48 h transfection with P4HB, ERp57 and ERp72 with presence or absence of ascorbate after 24 h incubation. Quantification of median fluorescence intensity was measured using FlowJoTMv10 software. Results are mean ± SEM of 3 independent experiments. Statistical analysis was performed by using two-way ANOVA with Tukey’s multiple comparison test. (b) Immunoblotting of WT-GFP-collagen and KO-GFP-collagen cells post 48 h transfection with P4HB in presence or absence of ascorbate after 24 h incubation for indicative antibodies and approximate molecular weight. (c) Filter-trap assay performed on the WT, KO, WT-GFP-collagen, and KO-GFP-collagen U2OS cell post 48 h transfection with V5-P4HB in presence or absence of ascorbate after 24 h incubation against collagen antibody. (d) Immunofluorescence on WT, KO, WT-GFP-collagen, and KO-GFP-collagen U2OS cell post 48 h transfection with V5-P4HB in presence or absence of ascorbate after 24 h incubation probed for GFP-collagen, KDEL, V5 and Dapi.

### Correlation between IRE1 signaling and P4HB levels in human samples from NASH patients

To explore the clinical relevance of our findings, we examined gene expression databases of liver samples from NASH patients. Previous studies have shown that the levels of XBP1s itself or that of its target genes predicts the survival of cancer patients [63, 64]. Thus, we divided RNA sequencing data from 155 NASH patients (human discovery cohort) into 2 groups, a high XBP1 expression group (77/155, high XBP1) and a low XBP1 expression group (77/155, low XBP1) (Figure 8a) [65, 66]. Remarkably, patients with low XBP1 gene expression signature pattern exhibited reduced P4HB levels, whereas the high XBP1 group had high P4HB mRNA levels (Figure 8a). In contrast, the levels of mRNA coding for α subunit of prolyl-4-hydroxylase P4HA1 were similar in the two groups differing in XBP1 expression (Figure 8a). In addition, the analysis of promoter DNA sequences of different isoenzymes of the prolyl-4-hydroxylase in different species revealed conserved core binding sequence of GCCACGT site for XBP1 (Figure S7a and S8). P4HB1 1 was the only isoenzyme with such a conserved core XBP1 binding site (Figure S7a) [67].

**Figure 8.**
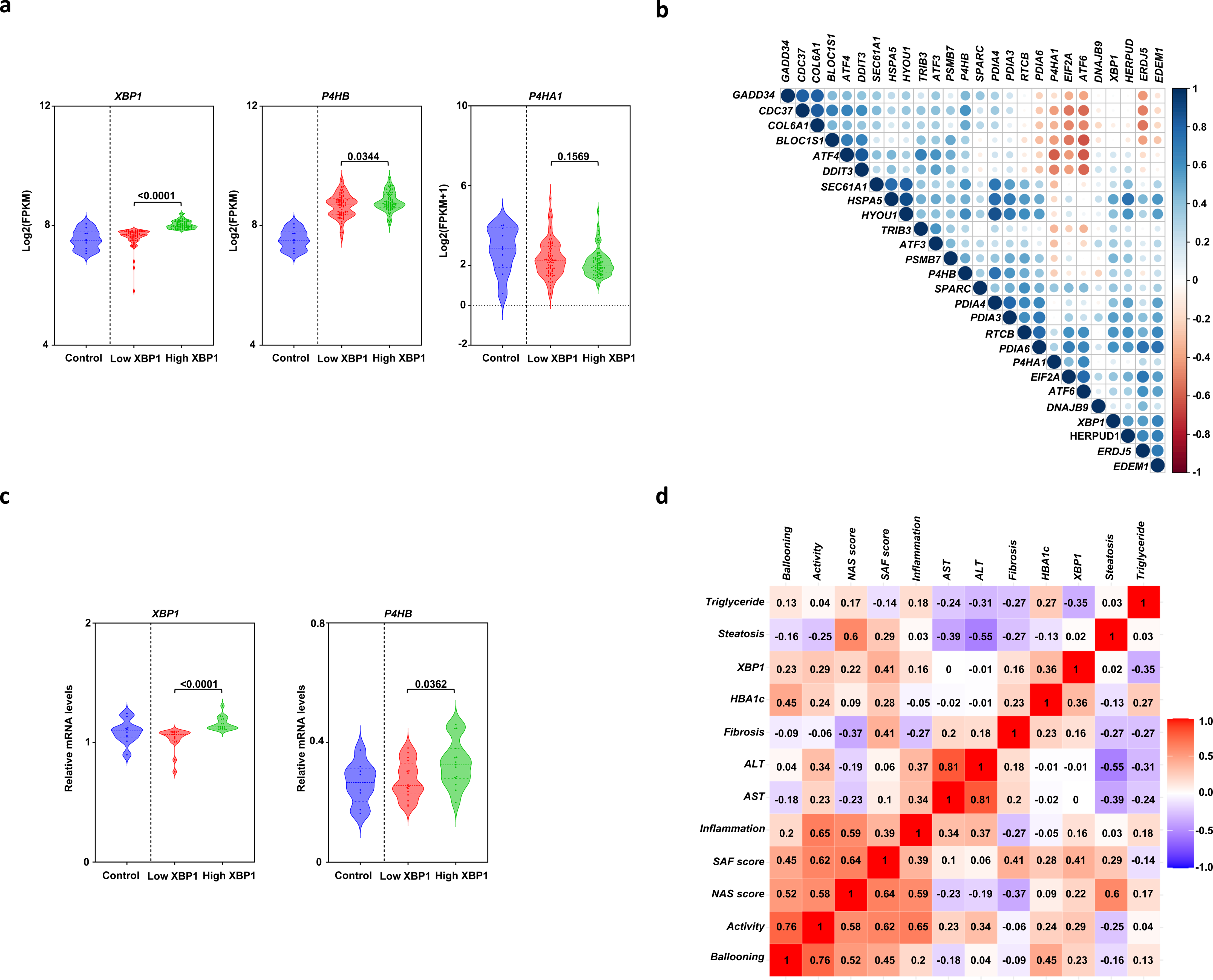
Correlation between XBP1 levels, P4HB expression and NASH in human patients. (a) The RNA seq data from 155 NASH patients were divided into high XBP1 and low XBP1 groups based on the quartile 2 (Q2) mean value for *XBP1*. Values above Q2 were called high XBP1(77, green) and values below Q2 were called low XBP1(77, red) of fragments per kilobase of exon per million mapped fragments (FPKM) of XBP1, P4HB and P4HA1 are shown 10 control samples are shown (blue). Statistical analysis was performed using one-way ANOVA with Tukey’s multiple comparison test. (b) Heatmap representing the Pearson Comparison of different UPR genes in RNA seq data of 155 NASH (c) qPCR of XBP1 in NAFLD human liver tissue samples of 10 controls and 36 NASH were divided into high XBP1 and low XBP1 groups based on the quartile 2 (Q2) mean value for XBP1. Values above Q2 were called high XBP1(16, green) and values below Q2 were called low XBP1(17, red). Relative mRNA of XBP1 and P4HB are shown. 10 control samples are shown (blue). Statistical analysis was performed using an unpaired t test with Welch’s correction. (d) Heatmap representing the Pearson Comparison of XBP1 and various clinical features of 36 NASH patients.

We also evaluated the levels of known UPR-target genes in the cluster of patients showing low and high XBP1 expression and observed a positive correlation (Figure S7b). Next, we evaluated the relation between XBP1 and P4HB (but not P4H1) in all NASH patients and detected a positive correlation (r= 0.31 and *p* < 0.0001) (Figure 8b). However, P4HA1 did not show any correlation with XBP1 in NASH human samples (Figure 8b). Analysis of expression levels of genes associated with hepatic fibrogenesis indicated a positive correlation between XBP1 and COL1A2, DCN and FBN (Figure S7c), important genes encoding ECM components [65, 68]. Further, levels of CTGF, a key ECM modulating protein, also showed positive correlation with XBP1 (Figure S7c). To validate our findings, we assessed a second cohort of NAFLD human liver tissue (French cohort), including 10 controls, 29 steatosis, 36 NASH cases. Remarkably, a direct correlation was again detected between XBP1s and P4HB levels in NASH liver samples (Figure 8c). Finally, we assessed the possible correlation between XBP1 levels and clinical data in this set of NASH patients. Remarkably, we observed a positive correlation between various clinical features and found positive correlation between XBP1 levels and SAF (steatosis, activity, and fibrosis)-defined score of activity (*r*=0.417 and *p*=0.052) (Figure 8d). XBP1 levels in NASH samples also correlated with hepatic ballooning, lobular inflammation, and augmenting fibrosis (Figure 8d). Overall, these results indicate that the levels of P4HB in the liver of patients with NASH significantly correlated with IRE1/XBP1 signaling and with disease prognosis, suggesting a role of the IRE1/P4HB axis in liver disease.

## Discussion

Collagen is the main ECM component in all tissue, representing the most abundant animal protein. Its abnormal deposition plays essential roles in the pathogenesis and progression of various disease states including fibrosis and cirrhosis [69, 70]. Collagen biosynthesis is a highly complex process because this protein undergoes extensive post translational modifications within the ER lumen before attaining its characteristics triple helical structure. This triple helix formation and its subsequent trafficking from the ER to the Golgi apparatus are critical for the secretion of stable collagen into the extracellular space. Collagen maturation critically involves several ER resident proteins like prolyl hydroxylases, BiP, HSP47 and PDIs [17, 71]. Prolyl 4-hydroxylase is an ascorbate-dependent tetramer enzyme composed by two α subunits (for which 3 isoforms exist: P4HA1, P4HA2, and P4HA3) and two conserved β subunits (P4HB or PDIA1) [72, 73]. P4HB is one of the abundant soluble proteins in the ER lumen and has multiple functions as reductase, oxidase, and isomerase, as well as a molecular chaperone [61, 74, 75], Prolyl 4-hydroxylase plays a central role in collagen synthesis because it catalyzes the hydroxylation of the C4 residue of proline yielding 4-hydroxyproline. This posttranslational modification in pre-pro-collagen is required for stable triple helix formation [76]. Thus, proteostasis within the ER is key for synthesis of stable triple helix collagen and alterations to this process will eventually lead to the production of unstable misfolded collagen.

IRE1 signaling is central for maintenance of tissue homeostasis and its deregulation due to chronic ER stress contributes to a variety of pathological conditions including hepatic steatosis, NASH, and fibrosis [77]. Here we employed distinct mouse models and disease paradigms, coupled to unbiased proteomic profiling, to ascertain the impact of IRE1 on liver disease models. We found altered expression of collagen-containing extracellular matrix in IRE1-defficient livers, indicating that the activity of the pathway regulates collagen stability and secretion. These effects were associated with altered expression of different factors involved in proteostasis control and collagen production, highlighting P4HB as one of the main downregulated gene in IRE1 deficient livers. Hepatic pathologies often progress with fibrosis, the adjustment of ER proteostasis according to the secretory need through the IRE1-P4HB axis appears to be an important adaptive mechanism to sustain homeostasis under continuous stress like steatosis, HFD, chronic alcohol consumption, acute toxic insults and even infection by hepatitis B or C [78, 79]. Further, collagens are the most abundant cargo of secretory output constituting about 12-17% of total mice protein and represents one of the most abundant proteins in the ECM deposition of liver fibrosis [80, 81]. In cirrhotic livers, collagen content increases four to seven-fold as compared to normal liver, and its production is closely coupled to the IRE1-P4HB pathway, possibly as a mechanism to enhance the secretory capacity of a cell and to avoid proteotoxicity due to the accumulation of misfolded collagen in HSCs [82, 83]. Based on our results, we propose that IRE1-P4HB pathway is essential to finetune the collagen folding capacity and secretory outcome in normal physiology and plays an important pathogenic role in excessive collagen deposition in the ECM. In addition to impacting liver toxicity and collagen deposition, ablation of IRE1 altered the differentiation process of HSCs, which may represent a parallel phenomenon contributing to the disease process. In this line, professional secretory immune cells lacking XBP1 display severe abnormalities in their development and function as initially reported in plasma B-cells [84] and dendritic cells [85]. XBP1 is important for B-cell differentiation before the engagement of a highly secretory demand through signaling downstream of the B cell receptor [86]. Consistent with this, analysis of the XBP1s gene regulatory network at a genomic scale revealed a key role of this UPR transcription factor in cell differentiation associated with regulation of Mist1, a master regulator of myoblast cell differentiation [67]. In the same way, XBP1 deficiency in gastric epithelial cells led to altered differentiation of digestive-enzyme–secreting zymogenic cells possibly due to Mist1 deregulation [87]. A novel function of XBP1 was also uncovered in the liver in the regulation of normal fatty acid synthesis [39]. It remains to be determined if XBP1 is part of a direct regulatory circuit controlling HSCs differentiation. In this line, it is interesting to note that XBP1 regulates the transcription of TANGO-1, a central component in the trafficking of collagen to the Golgi apparatus [16]. XBP1 also regulates key genes involved in fatty acid synthesis and targeting XBP1 in the liver triggered hypocholesterolemia and hypotriglyceridemia, secondary to a decreased production of lipids from the liver in the absence of hepatic steatosis [88]. Another study also showed that ablation of XBP1 ameliorates hepatosteatosis, liver damage, and hypercholesterolemia in an animal model of dyslipidemia [89], whereas other studies have shown that its expression regulates triglyceride-transfer protein downregulation to promote hepatosteatosis [40]. Here we report that collagen deposition is critically regulated by the IRE1 pathway in part through the control of P4HB expression. IRE1 is regulated by the assembly of a dynamic protein platform referred to as the UPRosome, which may control the stress threshold to engage the UPR, its temporal inactivation and the cross talk with other stress pathways [90]. In addition to BiP, other ER chaperones regulate IRE1 activation. Hsp47 binds to IRE1 and engages IRE1 signaling, involving the release of BiP from the complex. P4HB/ PDIA1 and PDIA6 also modulate IRE1 activation through the formation of a protein complex [91]. In addition, the phosphorylation of PDIA1 regulates IRE1 activity, constituting a post-translational response to maintain ER proteostasis and playing a vital role in protecting against ER stress-induced cell death. We speculate that the molecular connection identified here between IRE1 and P4HB/ PDIA1 may represent a central mechanism that adjusts the ER folding needs according to the collagen synthesis, meaning that exacerbated collagen expression stimulates IRE1 signals to improve the secretory capacity of the cell via an elevation of P4HB/PDIA1 [82]. Since XBP1s regulates TANGO-1, and the mRNA coding for collagen-6a is also a major RIDD substrate, sustaining an equilibrated pathway to fold collagen may represent a major “challenge” to the UPR. Indeed, studies in medaka fish indicated that the abnormal developmental phenotypes triggered by the genetic disruption of major UPR components are due to the accumulation of misfolded collagens [92]. Thus, a feedback loop exists between collagen biogenesis and adjustment of UPR signaling. The inhibition of HSCs activation and extracellular collagen deposition is the goal of anti-fibrotic therapies. We propose that pharmacological inhibition of IRE1 activation might represent a feasible strategy to protect against steatohepatitis and liver fibrosis.

## Materials and Methods

### Cell culture, cell lines and DNA constructs

U20S, Hepa 1-6 and HuH-7 were grown in Dulbecco’s modified Eagle’s medium–high glucose (DMEM, Gibco) supplemented with 10% fetal bovine serum (FBS), 100 units/ml penicillin, and 0.1 mg/ml streptomycin (Invitrogen) at 37°C under 5% CO2. For routine transfection, Effectene transfection reagent (Qiagen) was used as recommended by the manufacturer. CRISPR cells were generated using double nickase targeting IRE1 or scrambled as a control (control:sc-437281, IRE1 human:sc-400576-NIC and IRE1 mouse:sc-429758-NIC) (Santa Cruz). Minimum of four clones were screened from two different set of clones after probed for XBP1 mRNA splicing and the upregulation of BIP and CHOP which was induced by 250 ng/mL of tunicamycin (Tm) for 4, 8, 16 and 24 h (Tm kinetics) [93]. GFP-collagen stable were generated using constructed purchased from the addgene (addgene 110726).GFP-collagen expressing cells were made as described in [94].

### RNA isolation, RT-PCR, real-time PCR and Xbp1 mRNA splicing

Total RNA from cells and tissues were extracted with Trizol (Invitrogen, Carlsbad, CA, USA). cDNA was synthesized with random primers p(dN)6 (Roche) using SuperScript III Reverse Transcriptase (Invitrogen). Real-time quantitative PCR reactions using SYBRgreen fluorescent reagent and/or EvaGreen™ were achieved on the Stratagene Mx3000P system (Agilent Technologies, Santa Clara, CA 95051, USA) with specific set of primer (Supplementary Materials and Methods). The relative amounts of mRNAs were calculated from the values of comparative threshold cycle by using *Actin* mRNA as control. Xbp1 mRNA splicing assay have been previously described [20, 95, 96]. Semi-quantitative PCR was performed using specific primer pair (Supplementary Materials and Methods).

### Animal Study, CCl_4_ treatment and HDF model

All animal experiments were carried out according to the Guide for the Care and Use of Laboratory Animals of the National Institutes of Health. The protocol for use of animals in this study was approved by the Committee on the Ethics of Animal Experiments of the University of Chile (protocol number CICUA CBA 1015 FMUCH). Mice used were 2 to 4 months old and were age matched littermates housed at 23±2℃ with a humidity of 35±5% under a 12:12 h light-dark cycle (lights on at 7.00 am) with access to food and water *ad libitum.* Mice flox for RNase [42] or kinase domain [48] of *Ern1* were bred with Mx-Cre ([41] and deletion was carried out with 150 μg intraperitoneal poly I:C (invivogen) injection thrice after every two days [20, 93]. For induction of acute liver toxicity, single doses of 1.6 g/Kg of CCl_4_/olive oil (1:1 ratio) were injected intraperitoneally 2, 6, 12 and 24 h before the mice were sacrificed, and livers were harvested and immediately weighed. The same volume of the olive oil as a vehicle was injected intraperitoneally into the control mice. For chronic CCl_4_ treatment, 0.4 g/Kg of CCl_4_/olive oil (1:1 ratio) were intraperitoneally injected for 12 consecutive weeks with 3 doses per week on alternative days [43, 97]. The same volume of the olive oil as a vehicle was injected into the control mice. HFD experiments were conducted using 12-week-old mice with floxed littermates used as controls. Mice were randomly assigned for feeding with HFD (TestDiet, St.Louis, Missouri:58Y1) for 12 consecutive weeks [16, 38].

### Murine HSC isolation and treatment

Murine HSC isolations were performed in accordance with the IACUC protocol approved by the Indiana University School of Medicine (Protocol number 21025). mHSCs were isolated from IRE1^fl/fl^ mice (generously provided by Dr. Randy Kaufman) as previously described [48, 98, 99]. Briefly, mice were perfused via the portal vein with a combination of pronase and collagenase to digest the liver. HSCs were separated from other liver cell types using an accudens density gradient and cultured in DMEM supplemented with 10% fetal bovine serum (FBS), 100 units/ml penicillin, and 0.1 mg/ml streptomycin (Invitrogen) at 37°C under 5% CO2. mHSCs were infected with adenovirus encoding for AdCre-GFP or an AdLacZ-GFP control (University of Iowa Viral Vector Core). Following infection, mHSCs were treated with 2ng/mL TGF-β or vehicle for 24h. Conditioned media was harvested and concentrated using Amicon Ultra Centrifugal Filters (Millipore UFC503096). For protein analysis cells were washed 2X in cold PBS and lysed in RIPA buffer containing protease and phosphatase inhibitors, centrifuged to remove debris, and reduced by adding 4X Lamelli Buffer + β-mercaptoethanol and boiling samples. Protein from whole cell lysates and conditioned media were analyzed by running samples on SDS/polyacrylamide gel electrophoresis (4-15% Stain-free SDS-PAGE minigels) and transferred to PVDF membrane (Immun-Blot PVDF, Bio-Rad 1620177). Membranes were blocked in 3% BSA in Tris-buffered saline containing 1% Tween-20 (pH7.4), for 1hr at room temperature, incubated with primary antibody overnight at 4°C, followed by an appropriate secondary antibody, and developed using SuperSignal West Femto Maximum Sensitivity Substrate (Thermo Fisher Scientific) following manufacturer’s instructions. Chemiluminescence signal and protein ladder images were acquired using ChemiDoc Imaging System (BioRad). For qPCR analysis, isolated HSCs were washed 2X with cold PBS, and mRNA was isolated using a Qiagen RNeasy kit. mRNA was reverse transcribed using the First Strand cDNA Synthesis Kit (Applied Biosystems), and qPCR analysis was performed using a QuantStudio5 machine (Applied Biosystems).

### Liver staining and TUNEL analysis

CCl_4_ induced liver damage was analyzed with the help of DeadEnd^TM^ Fluorometric TUNEL system (Promega Corp G3250) according to given instructions[100]. Fluorescence from the liver sections were visualized by fluorescence microscopy and quantified using ImageJ [101]. Livers was fixed in 4% phosphate-buffered formaldehyde solution and embedded in paraffin.2 μm tissue sections were cut with sliding microtome for histology stained with either hematoxylin and eosin (H&E), Masson’s trichrome and Sirius Red staining [102–104].To remove paraffin of liver tissue slices, samples were treated with xylene twice for 5 minutes and subsequently rehydrated with absolute ethanol, 90% ethanol, 80% ethanol, 70% ethanol and washed with PBS 1X for 5 minutes. To antigen retrieval or epitope expose, slide was incubated 100 °C for 15 minutes in citrate buffer (Citric Acid diluted in H2O) 10 mM pH 6.0 in 0.05% PBS tween 20 or permeabilizated with Triton X-100 0.3%, BSA 1% in PBS 1X for 2 hours and the washed in PBS 1X 3×5 minutes both. Slides for immunohistochemistry were incubated in H2O2 3%, Methanol 10% diluted in PBS 1X 20 minutes, to block endogen peroxidase. Then slides or slices for immunofluorescence were incubated for 1 hour with blocking solution (BSA 1% in PBS 1X) at room temperature. Primary antibody incubation was made overnight at 4°C diluted in PBS 1X or blocking solution for immunohistochemistry or immunofluorescence, respectively. Secondary antibody was obtained from VECTASTAIN® Elite® ABC-HRP Kit, Peroxidase (Standard) PK-6100 or diluted in blocking solution and incubated for 1 hour at room temperature both, then washed with PBS 1X. Immunohistochemistry counterstaining was made employing DAB Peroxidase (HRP) Substrate Kit (with Nickel), 3,3’-diaminobenzidine by Vector Laboratories Inc SK-4100 and dehydrating with 70% ethanol 5 minutes, 80% ethanol 5 minutes, 90% ethanol 5 minutes, absolute ethanol 5 minutes and Xylene 2×5 minutes, then fixed with mounting solution. Instead, immunofluorescence was incubated with Hoesch 5 minutes, then fixed with ImmunoHistoMount™.

### Serum biochemistry and Liver measurements

Serum alanine transaminase and aspartate transaminase levels were measured using ALT (10.270.704) and AST (10.260.704) determination kits (DiaSys Diagnostic Systems GmbH, Holzheim, Germany). For acute liver toxicity experiments serum ALT and AST levels were measured with the help of VetLab (Laboratorio clinic veterinario using IFCC method). For hepatic triglycerides 50 mg of liver tissue was homogenized in PBS and subsequently mixed with 1.6 mL of chloroform. After centrifugation at 3000 rpm for 10 min at room temperature, residual volume was resuspended in absolute ethanol (Triton X-100). The triglyceride concentration was measured with the Serum Triglyceride determination Kit (Sigma) [27]. Hydroxyproline content was measured with Hydroxyproline Assay Kit (MAK008), 10 mg tissue was homogenized and hydrolyzed with 10 N NaOH at 120 °C for 3 hours. Lysate was cooled on ice and neutralized with 10 N HCl subsequently centrifuged. Supernatant was collected and quantified as per manufacturer’s instructions [105].

### Protein sample preparation, media protein precipitation, immunoblotting, and filter trap assay

Cells were harvested, resuspended in ice-cold PBS, and centrifuged (300 g, 5 min, 4℃) followed by a wash with ice-cold PBS. Cells were lysed in RIPA buffer (89901 PierceTM) containing protease inhibitor cocktail (Roche 04693124001) and PhosSTOP (Roche 04906837001) and kept frozen in −80℃ until analysis. Liver samples were homogenized in RIPA buffer containing protease inhibitor cocktail (Roche 04693124001) and PhosSTOP (Roche 04906837001) and sonicated (QSonica, Q125) for 15 s. Media protein was collected and centrifuged at 10,000 rpm for 10 min and precipitated using 1:1V/V ratio of cold acetone (−20℃). Tubes were vortexed and kept overnight at −80℃ followed by 15,000 rpm for 60 min. Supernatant was decant and pellet was dissolved in in RIPA buffer containing protease inhibitor cocktail (Roche 04693124001) and PhosSTOP (Roche 04906837001). Protein concentration was determined by micro-BCA assay (Pierce). Lysates were denatured using 4× Lamelli buffer (NuPage NP0008 novex) with beta-mercaptoethanol (Sigma M7154) and boiled. 20-50 μg of total protein was resolved using SDS/polyacrylamide gel electrophoresis (8-12% SDS-PAGE minigels) and transferred to PVDF membrane (Immobilon ISEQ00010). Membranes were blocked using PBS, 0.1% Tween20 (PBST) containing 5% milk for 60 min at room temperature incubated with primary antibodies overnight, followed by an appropriate secondary antibody, and developed using the ECL method (Thermo Fisher Scientific) following manufacturer’s instructions. Chemiluminescence signal and protein ladder images were acquired using ChemiDoc Imaging System (BioRad).

Filter trap assays were performed by sonication of protein fractions treated with 100 mM beta-mercaptoethanol or deionized water for 30 min on ice and then diluted in PBS with 1% SDS with final protein concentration of 0.25 μg/ μL to avoid clogging of the membrane pores. The protein samples were vacuum filtered through a 0.22 μm pore size cellulose acetate membrane. Membrane was then washed once with PBS-SDS and twice with PBS-T for 5 min at RT each and blocked with 5% non-fat dry milk in PBS and followed by routine immunoblotting procedure with different Anti-GAPDH (2118s, CST), Anti-collagen (1310-01, Southern Biotech), Anti-P4HB (ab137110, Abcam), HSP90 (SC-13119, Santa Cruz Biotechnology), Anti-V5 (R960-25, Invitrogen), Anti-PDI1 (SC-20132, Santa Cruz Biotechnology), Anti-PDI1 (Ab2792, Abcam), Anti-IRE1 (3294s, CST), Anti-Bip (Ab21685, Abcam), Anti-ERp72 (ADI-SPS-720-D, Enzo), Anti-ERp57 (SC-28823, Santa Cruz Biotechnology), Anti-KDEL (ADI-SPA-827, Enzo), Anti-P62 (ab56416, Abcam), Anti-Hsp47 (Ab109117, Abcam), Anti-Calnexin (ADI-SPA-860-F, Enzo), Anti-Calreticulin (ADI-SPA-600-F, Enzo), Anti-ERO1L (NB100-2525, Novus Biologicals), Anti-P62/SQTM1 (SC-25575, Santa Cruz Biotechnology), Anti-LC3 (4108s, CST), Anti-beta-Actin (5125S, CST), Anti-Collagen (Ab260043, Abcam), Anti-Collagen A1 (NB-600-450, Novus Biologicals), Anti-Poly-UB (BML-PW8805, Enzo), Anti-αSMA (ab5694, Abcam), Anti-P4502E1 (ab28146, Abcam) antibodies [106].

### Immunoprecipitation assays

WT or IRE1KO U20S cells stably expressing Collagen-GFP were grown in 10 cm-plate. Cell lysates were prepared for immunoprecipitation as described above for HEK cells. As a control, to eliminate non-specific binding, experiments were performed in parallel in WT U2OS cells. After 48 h, protein extracts were prepared in lysis buffer (0.5% NP-40, 150 mM NaCL, 150 mM KCl, 50 mM Tris pH 7.6, 5% glycerol, and protease/phosphate inhibitors). Immunoprecipitation assays were performed as previously described [96]. In brief, to immunoprecipitated GFP-tagged Collagen, 600 ug of protein extracts were incubated with mouse anti-GFP antibody (SCBT) for 2 h at 4 °C, followed by the incubation with 15 ul of SureBeads™ Protein G Magnetic Beads for 1 h at 4°C. Beads-Protein complexes were washed 4 times with Lysis Buffer and resuspended in 2X LDS sample buffer. Samples were heated for 5 min at 95 °C and resolved by SDS–PAGE 8% followed by immunoblotting analysis to detect P4HB presence.

### TEM and CLEM

U2OS were fixed with 2.5% glutaraldehyde (EM-grade), 0.01% picric acid and 0.1 M cacodylate buffer pH 7.4 for 2 h. Samples were stained with 2% osmium tetroxide in 0.1 M sodium cacodylate buffer for 2 h, then washed with water and stained with 1% uranyl acetate for 2 h. Stained samples were dehydrated on an acetone dilution series and embedded in Epon (Ted Pella). Finally,80-nm sections were cut, mounted on electron microscopy grids, and examined using a transmission electron microscope (Talos F200C G2; Philips) at Advanced Microscopy Unit of Pontificia Universidad Católica of Chile. TEM analysis was performed double blinded using ImageJ software.

For CLEM, the U2OS cells grown on gridded glass cover slips (MatTek) were imaged using a Zeiss Axio Vert widefield inverted microscope equipped with a Hamamatsu ORCA-Flash4.0 LT CMOS camera and a live cell incubator system (PeCon GmbH, Erbach, Germany) at 37°C and 5% CO_2_. Samples were imaged with transmission (to identify cell positions on the gridded glass) and fluorescence modes (Zeiss eGFP filter set 38, excitation 470/40 nm and emission 525/50 nm) using 10X Plan-Neofluar air objective (NA 0.3) and 63X Plan Apochromat oil objective (NA 1.4). Immediately after LM imaging, the cells were fixed with 2% glutaraldehyde (EM-grade) in 0.1 M cacodylate buffer, pH 7.4 supplemented with 2 mM CaCl_2_ and MgCl_2_ for 30 minutes at room temperature. The samples were then washed with buffer and post-fixed with 1% reduced osmium tetroxide in 0.1 M cacodylate buffer, pH 7.4 on ice, for 1 h. Prior infiltration into epoxy resin (hard, TAAB 812, Aldermaston, UK) the samples were dehydrated through increasing concentration of ethanol. After polymerization of the resin at 60°C for 16 h, the target area for trimming of the pyramid was identified according to the grid marks transferred to the block surface. 60-nm thick sections were cut using an ultramicrotome (Leica EM Ultracut UC7, Leica Mikrosysteme GmbH, Austria) and five sections, every third, were collected on Pioloform-coated single slot copper grids. TEM micrographs from the correlated cells were acquired using a transmission electron microscope (Jeol Ltd., Tokyo, Japan) operating at 80 kV with a bottom mounted CCD camera (Orius SC 1000B, AMETEK Gatan Inc., Pleasanton, CA).

### FACS, Bodipy staining and cell sorting

GFP-Collagen U2OS cells were grown in the 12 well plate. After 24 hours treatment with ascorbate for overnight. Media was removed and trypsinization as routine subsequently resuspended in the 500 μL of PBS followed by flow cytometry using FACScan and FACSCalibur systems (BD Biosciences). Data was analyzed with FlowJo™ v10 Software (BD Biosciences).

GFP-collagen expressing cells were grown in 150 mm culture dishes, trypsinized and about 10-15 million cells were resuspended in 1 mL PBS with 10% FBS and 2 mM EDTA. The sorting cells were collected on the bases of normalized GFP signal using BD FACSAria™ III Sorter in 1 mL pure filtered FBS, centrifuged and seed in a 6 well plate for further processing.

The day before imaging U2OS cells were seeded in 12 well plates. After 24 h the cells were washed 3 times with prewarmed Hank’s Balanced Salt Solution (HBSS, Thermo Fisher) and stained with 1 μM of Bodipy red ER-tracker™ (Thermo Fisher E34250). After 1 h of incubation at 37 °C the cells were washed 3 times with HBSS followed by trypsinization and signal was measured using FACScan and FACSCalibur systems (BD Biosciences). Data was analyzed with FlowJo™ v10 Software (BD Biosciences).

### Immunofluorescence

P4HB-V5, Collagen, GFP-collagen and KDEL were visualized using immunofluorescence. Cells were fixed with 4% paraformaldehyde for 20 min at room temperature and later permeabilized with 0.5% TritonX constituted in PBS 0.5% bovine serum albumin (BSA) for 20 min followed by blocking for 2 h with 10% FBS in PBS 0.5% BSA. This was followed by overnight incubation with corresponding antibodies at 4℃ and subsequently washed and probed with Alexa-conjugated secondary antibodies (Molecular Probes) for 1 h at 37℃ with 15 min incubation with Hoechst. Coverslips were mounted with Fluoromount G onto slides and visualized by confocal microscopy (Fluoview FV1000).

### Affymetrix gene array analysis

Differentially expressed genes in mouse liver of *Ern1*/*Ern1^ΔR^* or *Ern1*/*Ern1^ΔK^* were determined using Affymetrix A450 2.0 gene chips [43]. Pre-processing and Normalization of Affymetrix arrays using linear probe level models (PLM) (affyPLM-package) and custom chip definition files (BrainArray Version 22; EntrezID, Clariom_S_Mouse) Differentially expressed genes were defined by a cutoff of 1.5-fold over levels in control liver tissue (p-value <10-5, FDR adjusted).

### Quantitative proteomic analysis

Liver samples were homogenized in RIPA buffer (89901 Pierce^TM^) containing protease inhibitor cocktail (Roche 04693124001) and PhosSTOP (Roche 04906837001). 10 μg of protein in 100 μl of RIPA were washed by chloroform/methanol precipitation and subsequently processed for mass spectrometry as described [107]. Air-dried pellets were resuspended in 1% RapiGest SF (Waters) and reconstituted in 100 mM HEPES (pH 8.0). Reduction of proteins was achieved with 5 mM Tris(2-carboxyethyl)phosphine hydrochloride (Thermo Fisher) for 30 min and samples were subsequently alkylated in 10 mM iodoacetamide (Sigma Aldrich, St. Louis, MO) for 30 min at ambient temperature and protected from light. Then proteins were digested with 0.5 μg trypsin/LysC (Thermo Fisher) for 18 hr at 37°C. Individual digested samples were reacted with the appropriate TMTpro-NHS isobaric reagent (Thermo Fisher) in 40% (v/v) anhydrous acetonitrile and quenched with 0.4% ammonium bicarbonate for 1 h. Different TMT labelled samples were pooled and acidified with 5% formic acid. Acetonitrile was evaporated on a SpeedVac and debris was removed by centrifugation for 30 min at 21,000 x g. MuDPIT microcolumns were prepared as described [108]. LC-MS/MS analysis was performed using an Exploris 480 mass spectrometer equipped with an Ultimate 3000 RSLCnano system (Thermo Fisher). MuDPIT experiments were performed by 5 min sequential injections of 0, 10, 20, 30, …, 100% buffer C (500 mM ammonium acetate in buffer A) and a final step of 90% buffer C / 10% buffer B (20% water, 80% acetonitrile, 0.1% fomic acid, v/v/v) and each step followed by a gradient from buffer A (95% water, 5% acetonitrile, 0.1% formic acid) to buffer B. Electrospray was performed directly from the analytical column (100µm ID fused silica, 20cm length, with a laser-pulled tip, filled with Aqua 3µm 10 Å C18 resin (Phenomenex) by applying a voltage of 2.2 kV with an inlet capillary temperature of 275°C. Data-dependent acquisition of MS/MS spectra was performed with the following settings: eluted peptides were scanned from 400 to 1600 m/z with a resolution of 120,000 and the mass spectrometer in top-speed data dependent acquisition mode. Peaks from each full scan were fragmented for 3 sec by HCD using a normalized collision energy of 32, a resolution of 45,000, 0.4 m/z isolation window, automatic injection time, AGC target of 200%, and scanned with a first mass starting at 110 m/z. Dynamic exclusion was set to 45 sec and 20 ppm. Peptide identification and protein quantification was performed using Proteome Discoverer 2.4 (Thermo Fisher). Normalization of TMT reporter ion intensities was carried out based on total peptide abundance in each channel, and subsequently, TMT intensity ratios for each identified protein were calculated between sample groups: *Ern1*/*Ern1^ΔR^* olive oil/CCl_4_ Acute treated (n=4), *Ern1*/*Ern1^ΔR^* olive oil/CCl_4_ chronic treated (n=4) and *Ern1*/*Ern1^ΔR^* HFD (n=7 for WT, one sample was not included because the sample in this channel showed lower abundance for unnormalized peptide; n=8 for KO). Significance assessed by a two-tailed unpaired t-test using the FDR approach [109] and Q=5% in Graphpad Prism. Enrichment analysis of most significant alterations between groups was performed in EnrichR platform or STING using gene ontology (GO) database [110–113]. Protein–protein interaction network was generated in STRING v.11[114].

### Metabolomics studies

Metabolic studies were performed in liver tissue samples as previously described.[115]. Approximately 30 mg of tissue for each condition was first weighed and solubilized in 1.5 mL polypropylene microcentrifuge tubes with ceramic beads using 1 mL of cold lysate buffer (methanol:water:chloroform, 9:1:1, 20°C). They were then homogenized three times for 20 s at 5,500 rpm. using a Precellys 24 tissue homogenizer (Bertin Technologies) followed by centrifugation (10 min at 15,000 g, 4°C). The upper phase of the supernatant was divided into two parts: the first 150 µl was used for the gas chromatography coupled to mass spectrometry (GC–MS) by vial injection, the other 250 µl was used for the ultrahigh pressure liquid chromatography coupled to mass spectrometry (UHPLC–MS). The GC–MS aliquots (150 µl) were evaporated and dried, and 50 µL of methoxyamine (20 mg ml^−1^ in pyridine) were added, after which the aliquots were stored at room temperature in the dark for 16 h. The following day, 80 µl of MSTFA was added and the final derivatization was performed at 40 °C for 30 min. Samples were then directly injected directlyinto GC–MS. For the LC–MS aliquots, the collected supernatant was evaporated into microcentrifuge tubes at 40 °C in a pneumatically assisted concentrator (Techne DB3). The LC–MS dried extracts were solubilized with 450 µl of MilliQ water and aliquoted into three microcentrifuge tubes (100 µl) for each LC method and one microcentrifuge tube as backup. Aliquots for analysis were transferred into LC vials and injected into UHPLC–MS or kept at −80 °C until injection. Subsequently, manual verification and quality control protocols were performed. An extended version of these methods is available upon request. The analysis of relevant metabolic pathways that were altered was obtained by analyzing the significantly altered metabolites with MetaboAnalyst 5.0[116].

### RNA-Seq analysis

RNA sequencing data used in this study was availed from NCBI GEO repository (GSE135251). The cohort comprised of 206 snap-frozen biopsy samples from patients diagnosed with NAFLD in France, Germany, Italy, or the United Kingdom and 10 healthy obese control cases without any biochemical or histological evidence of NAFLD from patients undergoing bariatric surgery in France. The cohort was stratified into according to assigned group in the paper [65]: NAFL (51), NASH F0 F1 (34), NASH F2 (53), NASH F3 (54), NASH F4 (14). Data were tested for normality using the Kolmogorov-Smirnov/ Shapiro-Wilk normality test using IBM SPSS software (IBM Corp). The RNA seq data from 155 NASH patients were divided into high XBP1 and low XBP1 groups based on the quartile 2 (Q2) mean value. Values above Q2 were called high XBP1 (77) and values below Q2 were called low XBP1 (77). Data were tested for normality using the Kolmogorov-Smirnov and significance was calculated using nonparametric Kruskal-Wallis test [117].

## Statistical analysis

Statistics were performed using Graphpad Prism 9.4.1 (GraphPad Software). Data were compared using One-way ANOVA or Two-way ANOVA for unpaired groups followed by multiple comparison post-test to compare more than two groups as stated in each figure. Student’s t-test was performed for unpaired group comparison between two groups. For proteomic experiment, statistical analysis was performed using multiple t-test with two-stage step-up method of Benjamini, Krieger and Yekutieli, with a False Discovery Rate of 5% in Graphpad prism 9.0. Post hoc testing for comparisons was performed using Turkey’s multiple comparison test.

## Author contributions

YH and CH designed the study. YH, HR, VAGL, JD, GT, MM, PP, SD, FA, RB, MH, XCD and ABG performed the experiments. BBM, JLM, AC, HV, LP, EJ, JGH, PG, GK supervised the experiments. YH analyzed the data and generated the figures for the manuscript. YH and CH wrote the manuscript. All authors read and approved the final version of the manuscript.

## Acknowledgments

YH is funded by FONDECYT no. 3180427. This work was funded by U.S. Air Force Office of Scientific Research FA9550-21-1-0096, FONDAP program 15150012, Department of Defense grant W81XWH2110960, ANID/FONDEF ID1ID22I10120, and ANID/NAM22I0057 and Swiss Consolidation Grant-The Leading House for the Latin American Region (CH). VAGL and LP are funded by NIGMS R35GM133552. BBM is funded by ANR “IRE1inNASH”. JLM is funded by NIDDK R03DK129326. XCD is funded by NIDDK DK120689, DK121925. MM is funded by a postdoctoral fellowship from ANID/FONDECYT No. 3200718. The authors acknowledge Euro-BioImaging (www.eurobioimaging.eu) for providing access to imaging technologies and services via the Finnish Advanced Microscopy Node (EMBI, Helsinki, Finland). EMBI is supported by Helsinki Institute of Life Science and Biocenter Finland.GK is supported by the Ligue contre le Cancer (équipe labellisée); Agence National de la Recherche (ANR) – Projets blancs; AMMICa US23/CNRS UMS3655; Association pour la recherche sur le cancer (ARC); Cancéropôle Ile-de France; European Research Council Advanced Investigator Grand “ICD-Cancer”, Fondation pour la Recherche Médicale (FRM); a donation by Elior; Equipex Onco-Pheno-Screen; European Joint Programme on Rare Diseases (EJPRD); European Research Council (ICD-Cancer), European Union Horizon 2020 Projects Oncobiome and Crimson; Fondation Carrefour; Institut National du Cancer (INCa); Institut Universitaire de France; LabEx Immuno-Oncology (ANR-18-IDEX-0001); a Cancer Research ASPIRE Award from the Mark Foundation; the RHU Immunolife; Seerave Foundation; SIRIC Stratified Oncology Cell DNA Repair and Tumor Immune Elimination (SOCRATE); and SIRIC Cancer Research and Personalized Medicine (CARPEM). This study contributes to the IdEx Université de Paris ANR-18-IDEX-0001.

## Supplementary Material

Supplementary data to this article can be found online at

## Supplementary figures

**Figure S1.**
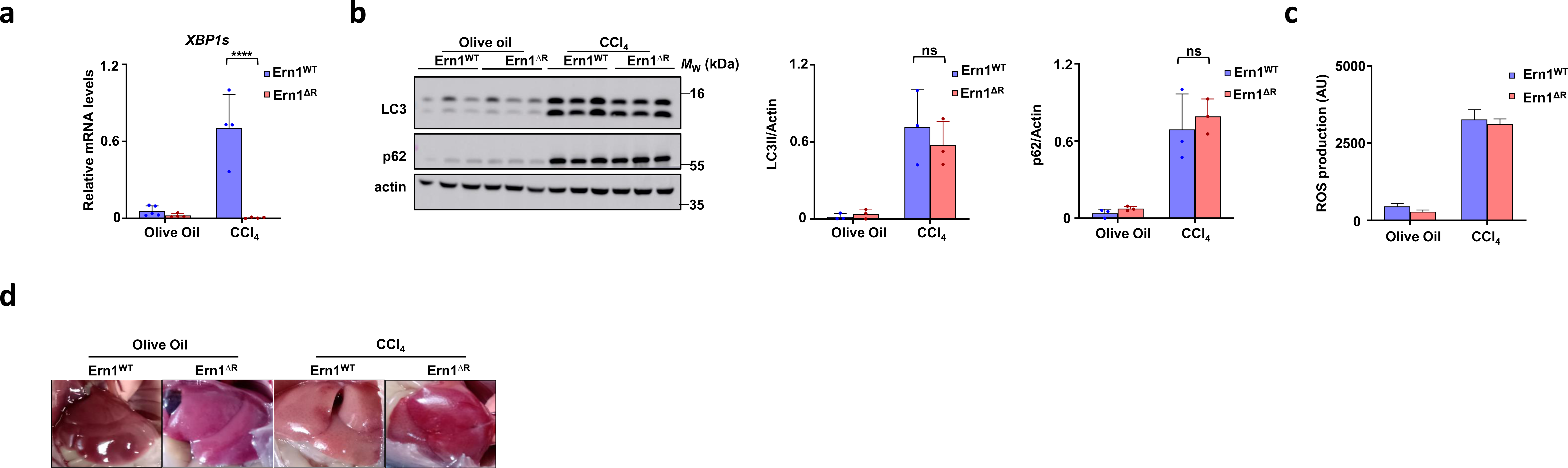
Comparable basal cellular outcomes due to *Ern1* deficiency. *Ern1* and *Ern1^ΔR^*age-matched mice littermates were treated with CCl_4_ for 24 h. (a) mRNA levels of *XBP1s* of *Ern1* and *Ern1^ΔR^* olive oil/CCl_4_ (1.6 g/Kg, i.p.) treated mice liver lysate after 24 h h (n=5 for *Ern1^WT^* olive oil and n=4 per group for rest. (b) Immunoblotting of liver extracts using the indicated antibodies and approximate molecular weight and bar graphs showing the corresponding relative protein quantification. Each lane represents an individual animal (n=3 per group). (c) Reactive oxygen species (ROS) of livers were measured (N1). (d) Representative images of liver appearance *in situ*. Statistically significant differences were determined by Two-Way ANOVA followed by Sidak’s multiple comparisons test unless mentioned.

**Figure S2.**
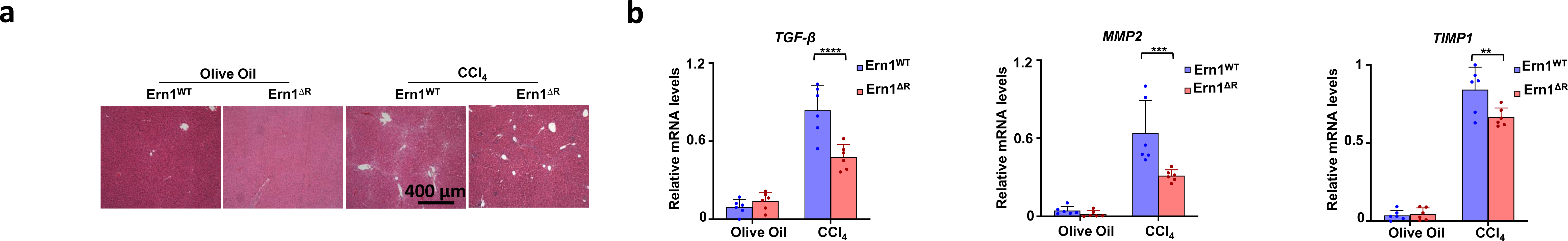
IRE1 deficiency suppresses the expression of fibrosis-related genes in mouse livers. (a) Representative H&E staining of liver. (b) mRNA levels of *TGF-β*, *MMP2* and *TIMP1* of *Ern1* and *Ern1^ΔR^* olive oil/CCl_4_ fibrotic mice model. Statistically significant differences were determined by Two-Way ANOVA followed by Sidak’s multiple comparisons test unless mentioned.

**Figure S3.**
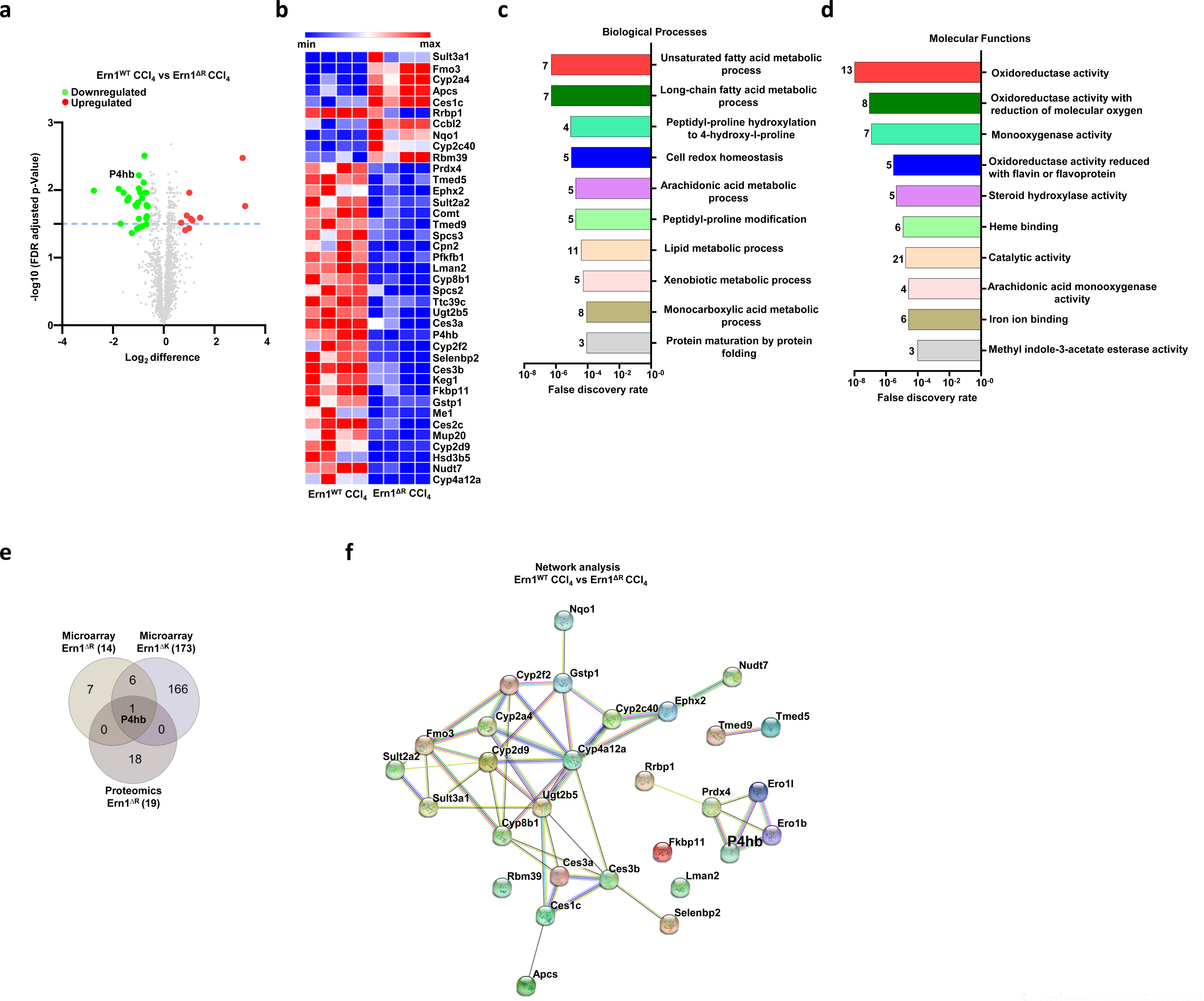
IRE1 deficiency alters proteostasis pathways in the liver. (a) Volcano plot of proteomic analysis of CCl_4_ treated *Ern1* and *Ern1^ΔR^* mice liver. The x-axis represents the logarithmic fold change levels in the *Ern1^ΔR^* mice liver relative to the *Ern1* mice liver and y-axis represents the negative decade logarithm of FDR adjusted P-values (n=4). Green dots indicate the downregulated proteins and red dots indicate the upregulation of proteins. Statistical analysis of unpaired t test with 5% False discovery rate using Two-stage step-up Benjamini. Krieger and Yekutiele method assuming individual variance for each row. Selected hits shown have q value ≤ 0.05 and P-value ≤ 0.05. Green dots indicate the downregulated proteins and red dots indicate the upregulation of proteins. (b) The heatmap represents the 29 downregulated proteins and 10 upregulated proteins from the *Ern1* and *Ern1^ΔR^* mice liver. Each lane represents individual mice and color intensity denotes the abundance of the protein. (c) Functional classification of proteomic hits according to the Gene Ontology (GO) annotation. Graph shows significantly enriched GO terms of biological processes. The number of Genes associated with each biological process is indicated in the parenthesis. (d) Functional classification of proteomic hits according to the Gene Ontology (GO) annotation. Graph shows significantly enriched GO terms of molecular functions. The number of Genes associated with each biological process is indicated in the parenthesis. (e) Vien diagram representing the common hit from microarray analysis of *Ern1^ΔR^* mice liver, microarray analysis of *Ern1^ΔR^* mice liver and proteomic analysis of *Ern1^ΔR^* mice liver. (f) Network analysis of P4HB interaction with identified genes related to protein maturation and collagen synthesis and secretion. The nodes represent the proteins and the edges interactions. The thickness of the edges indicates the extent of interactions, which does not necessarily mean physical binding (Ero1 and Ero1b included to show proximity).

**Figure S4.**
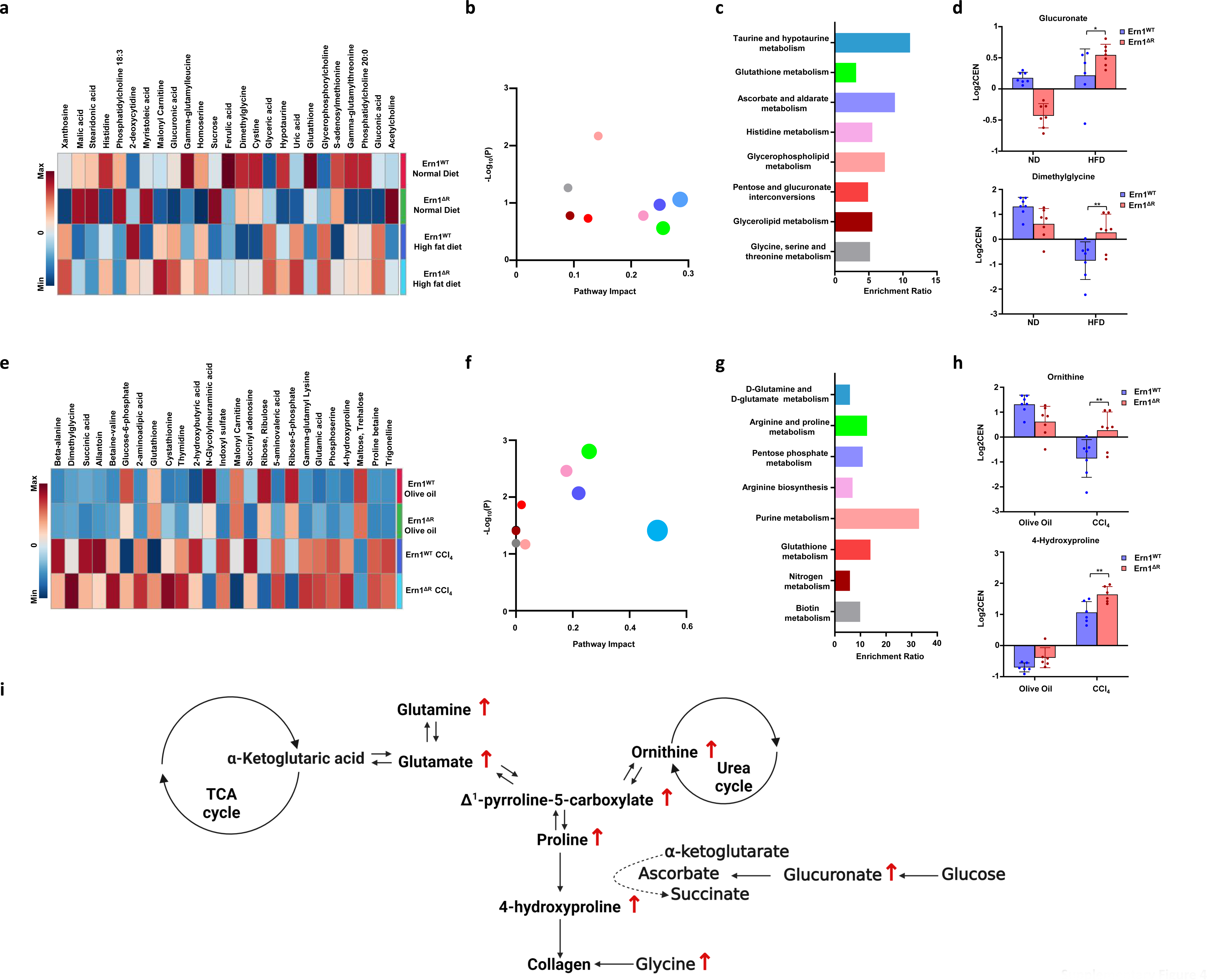
Metabolomic profiling of IRE1 deficient livers. (a) Heat maps of different metabolites of high fat diet fed *Ern1* and *Ern1^ΔR^* mice liver samples indicating the top 25 metabolite hits (n=6 or 7 per group). The color intensity is representative of the average of each group. (b) Pathway analysis showing the impact of top 8 the pathways. P values were calculated using hypergeometric test (c) Pathway enrichment indicates the ratio of number of hits in top 8 affected pathways to expected value of hits. (d) The affected metabolites in collagen metabolism i.e., the Glucuronate and Dimethylglycine in different groups are shown as scatter dot plot. P value is calculated using 2way ANOVA. (e) Heat maps of different metabolites of CCl_4_ treated *Ern1* and *Ern1^ΔR^* mice liver samples indicating the top 25 metabolite hits (n=6 or 7 per group). The color intensity is representative of the average of each group. (f) Pathway analysis showing the impact of top 8 pathways. *P* values were calculated using hypergeometric test (g) Pathway enrichment indicates the ratio of number of hits in top 8 affected pathways to expected value of hits. (h) The affected metabolites in collagen metabolism i.e., the ornithine and 4-hydroxyproline in different groups are shown as scatter dot plot. P value is calculated using 2-way ANOVA.

**Figure S5.**
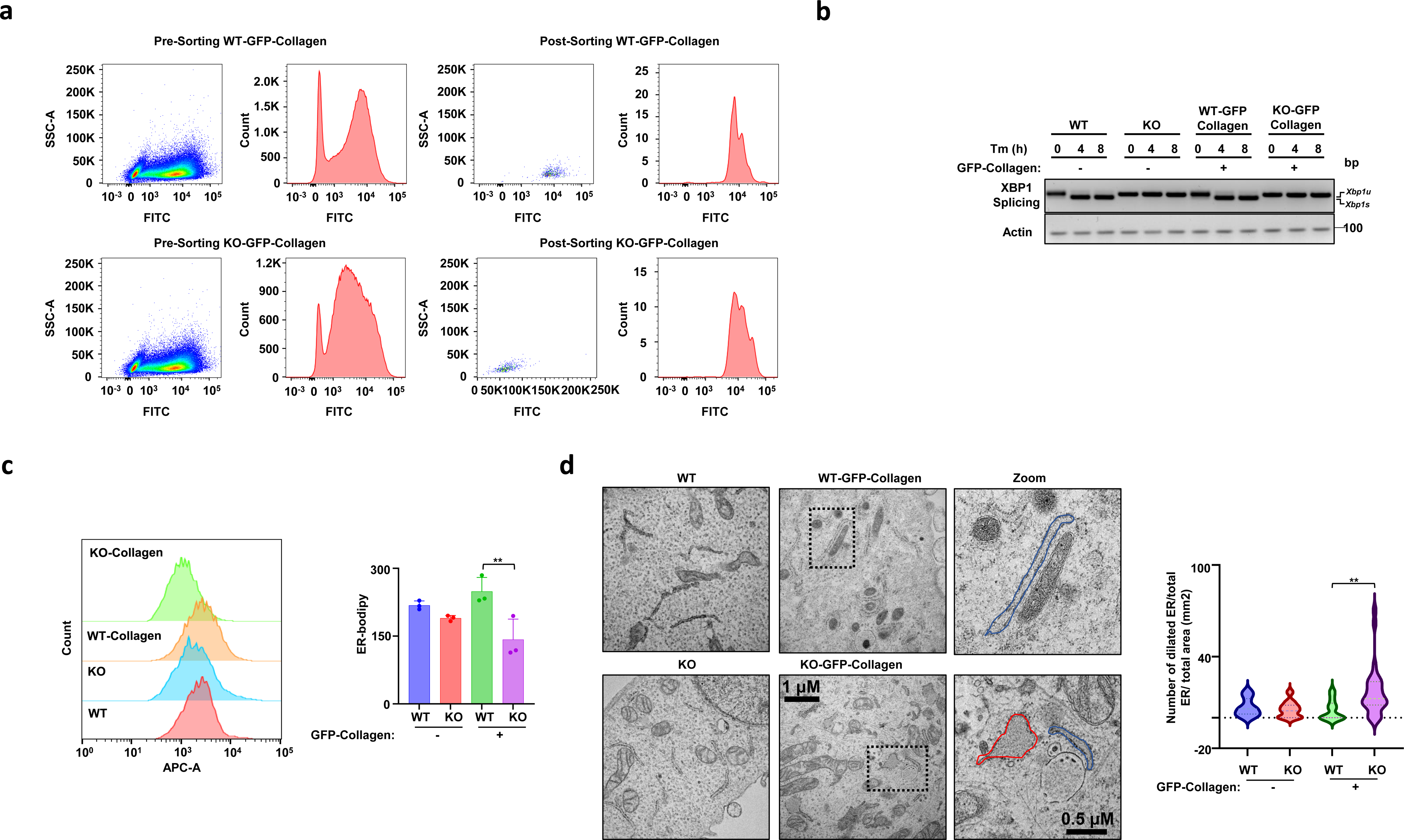
Control experiments and altered ER/Golgi content in IRE1 null cells. **(a)** (a) Represents the sorting strategy for selecting cells based on normalized GFP signal. **(b)** Regular splicing assay for Xbp1 mRNA for WT, KO, WT-GFP-collagen, and KO-GFP-collagen after 4 and 8 h of Tm treatment. **(c)** Representative fluorescence of Bodipy red ER-tracker indicating the extent of stained red stained ER. Data was analyzed using FlowJoTMv10 software. Quantification represents the median fluorescence intensity. Results are mean ± SEM of 3 independent experiments. Statistical analysis was performed by using one-way ANOVA with Tukey’s multiple comparison test. (d) Representative TEM images of CCT, KO, CCT-GFP-collagen, and KO-GFP-collagen U2OS cells showing the morphology and extent of ER (scale bar 1 μm). Quantification represents the area of ER (μm^2^) including above Q1 and below Q3. Data represents mean ±SEM from one experiment (n=23 CCT cells, n=55 KO cells, n=41 CCT- collagen cells and n=65 KO-collagen cells). Statistical analysis was performed by using one-way ANOVA with Tukey’s multiple comparison test.

**Figure S6.**
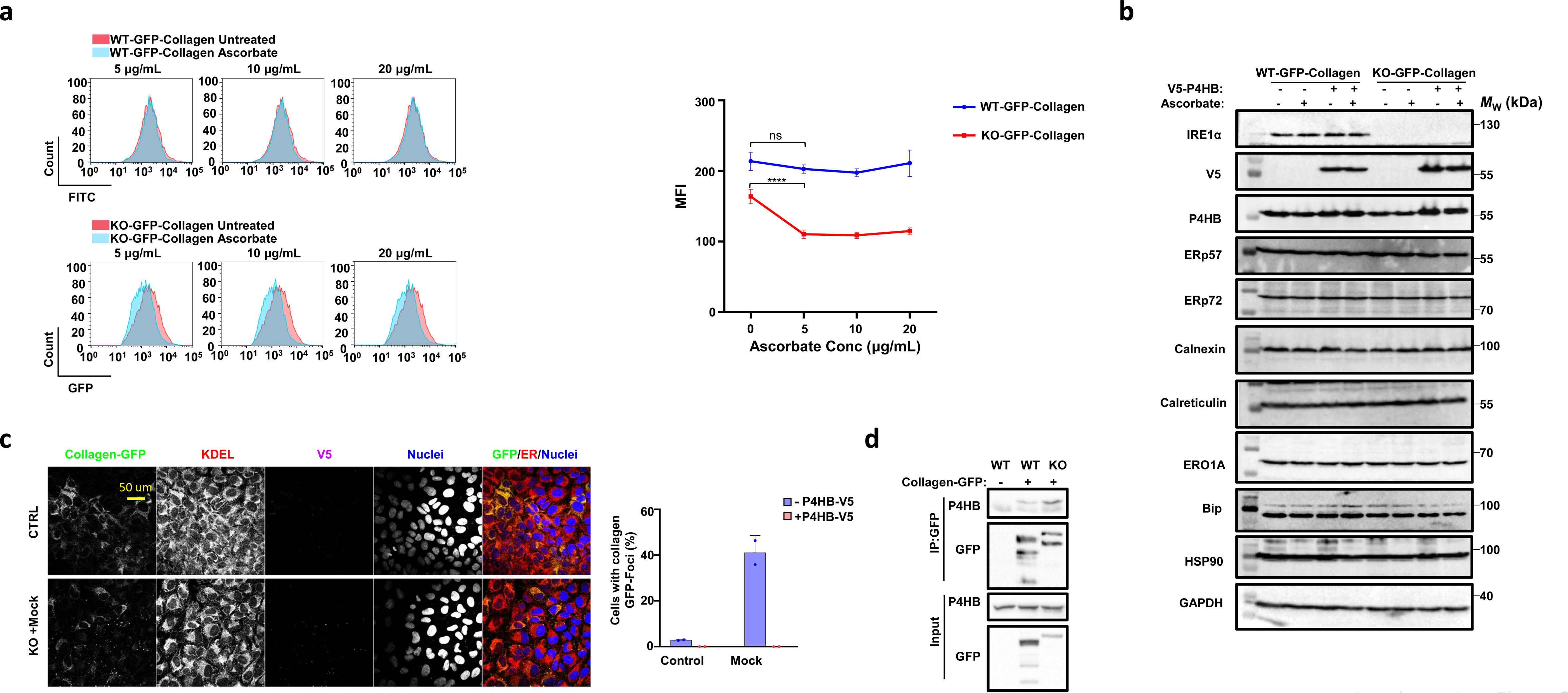
IRE1 deficiency does not affect all ER resident proteins or foldases. (a) Representative histograms of GFP-collagen fluorescence in presence of indicating ascorbate concentrations for 24 h. Quantification of median fluorescence intensity were measured using FlowJo^TM^v10 software. Results are mean ± SEM of 4 independent experiments. Statistical analysis was performed by using two-way ANOVA with Tukey’s multiple comparison test. (b) Immunoblotting of WT-GFP-collagen and KO-GFP-collagen cells post 48 h transfection with P4HB in presence or absence of ascorbate after 24 h incubation for indicative antibodies and approximate molecular weight. (c) Immunofluorescence on WT, KO, WT-GFP-collagen, and KO-GFP-collagen U2OS cell post 48 h transfection with V5-P4HB in presence or absence of ascorbate after 24 h incubation probed for GFP-collagen, KDEL, V5 and Dapi. (d) Co-immunoprecipitation of P4HB with WT-GFP-collagen or KO-GFP-collagen U2OS cells. Total extracts and IP were analyzed by Western blot using specific antibodies for P4HB and GFP. WT U2OS cells were used as control of immunoprecipitation.

**Figure S7.**
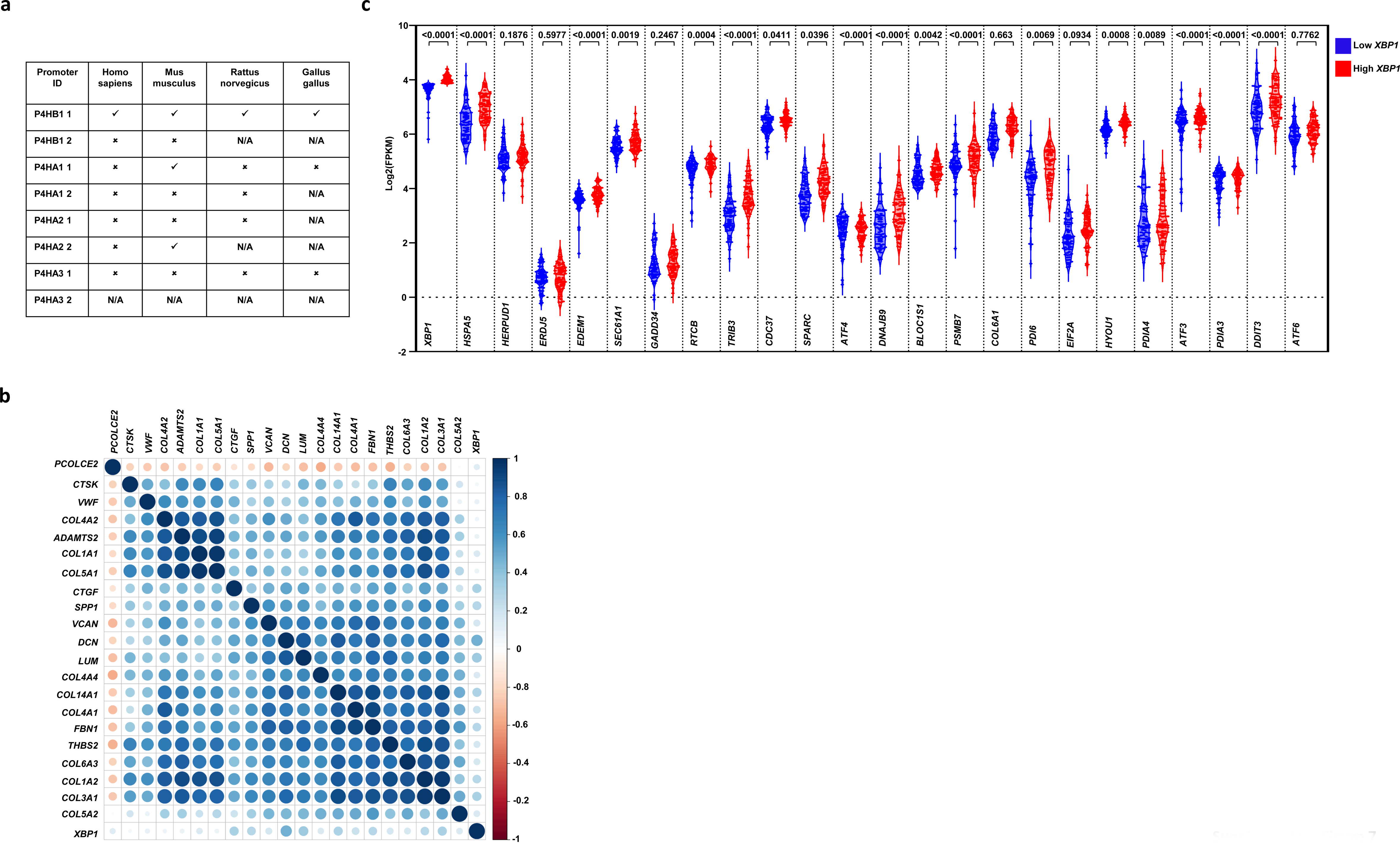
Correlation between XBP1 levels and fibrogenesis signaling signature in human NASH. (a) Represents the confirmation of conserved core sequence of GCCACGT site for XBP1 in the promoter sequence from −50 to −500 of different isoenzymes prolyl 4-hydroxylase in *Homo sapiens*, *Mus musculus*, *Rattus norvegicus* and *Gallus gallus.* (b) The RNA seq data from 155 NASH patients were divided into high XBP1 and low XBP1 groups based on the quartile 2 (Q2) mean value for *XBP1*. Values above Q2 were called high XBP1(77, green) and values below Q2 were called low XBP1(77, red) and logarithm of fragments per kilobase of exon per million mapped fragments (FPKM) of different UPR genes are shown. 10 control samples are shown (blue). Statistical analysis was performed using one-way ANOVA with Tukey’s multiple comparison test (c) Heatmap representing the Pearson Comparison of different genes involved in fibrogenesis in RNA seq data of 155 NASH.

**Figure S8.**
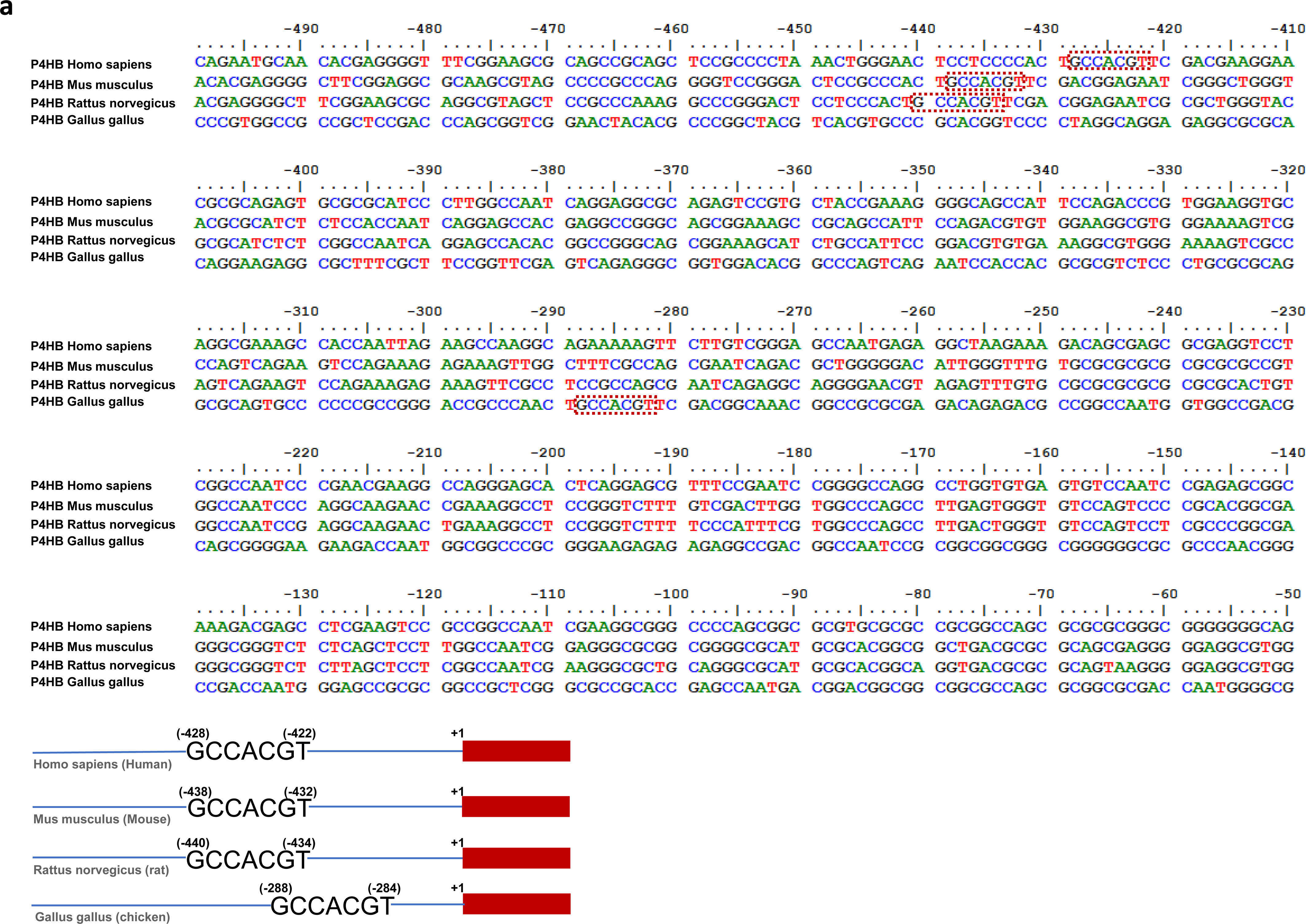
Sequences of P4HB promoter regions in different organisms showing the conserved core sequence of GCCACGT site for XBP1. Represents the promoter sequence from −50 to −500 of P4HB in *Homo sapiens*, *Mus musculus*, *Rattus norvegicus* and *Gallus gallus* showing the conserved core sequence of GCCACGT site for XBP1.

## References

1. Ishida, Y. and K. Nagata, Hsp47 as a collagen-specific molecular chaperone. Methods Enzymol, 2011. 499: p. 167–82.

2. Di Lullo, G.A., et al., Mapping the ligand-binding sites and disease-associated mutations on the most abundant protein in the human, type I collagen. J Biol Chem, 2002. 277(6): p. 4223–31.

3. Forlino, A. and J.C. Marini, Osteogenesis imperfecta. Lancet, 2016. 387(10028): p. 1657–71.

4. Jobling, R., et al., The collagenopathies: review of clinical phenotypes and molecular correlations. Curr Rheumatol Rep, 2014. 16(1): p. 394.

5. Murata, K., et al., Changes of collagen types at various stages of human liver cirrhosis. Hepatogastroenterology, 1984. 31(4): p. 158–61.

6. Rojkind, M. and R. Perez-Tamayo, Liver fibrosis. Int Rev Connect Tissue Res, 1983. 10: p. 333–93.

7. Blanck, T.J.J. and B. Peterkofsky, The stimulation of collagen secretion by ascorbate as a result of increased proline hydroxylation in chick embryo fibroblasts. Archives of Biochemistry and Biophysics, 1975. 171(1): p. 259–267.

8. Goldberg, B., E.H. Epstein, Jr., and C.J. Sherr, Precursors of collagen secreted by cultured human fibroblasts. Proc Natl Acad Sci U S A, 1972. 69(12): p. 3655–9.

9. Canty, E.G. and K.E. Kadler, Procollagen trafficking, processing and fibrillogenesis. J Cell Sci, 2005. 118(Pt 7): p. 1341–53.

10. Gorres, K.L. and R.T. Raines, Prolyl 4-hydroxylase. Critical reviews in biochemistry and molecular biology, 2010. 45(2): p. 106–124.

11. Jimenez, S., M. Harsch, and J. Rosenbloom, Hydroxyproline stabilizes the triple helix of chick tendon collagen. Biochem Biophys Res Commun, 1973. 52(1): p. 106–14.

12. Mussini, E., J.J. Hutton, Jr., and S. Udenfriend, Collagen proline hydroxylase in wound healing, granuloma formation, scurvy, and growth. Science, 1967. 157(157): p. 927–9.

13. Pihlajaniemi, T., R. Myllylä, and K.I. Kivirikko, Prolyl 4-hydroxylase and its role in collagen synthesis. Journal of Hepatology, 1991. 13: p. S2–S7.

14. Satoh, M., et al., Intracellular interaction of collagen-specific stress protein HSP47 with newly synthesized procollagen. J Cell Biol, 1996. 133(2): p. 469–83.

15. Ito, S. and K. Nagata, Biology of Hsp47 (Serpin H1), a collagen-specific molecular chaperone. Semin Cell Dev Biol, 2017. 62: p. 142–151.

16. Maiers, J.L., et al., The unfolded protein response mediates fibrogenesis and collagen I secretion through regulating TANGO1 in mice. Hepatology, 2017. 65(3): p. 983–998.

17. Nagata, K., HSP47 as a collagen-specific molecular chaperone: function and expression in normal mouse development. Semin Cell Dev Biol, 2003. 14(5): p. 275–82.

18. Bataller, R. and D.A. Brenner, Liver fibrosis. Journal of Clinical Investigation, 2005. 115(4): p. 1100–1100.

19. Hetz, C. and F.R. Papa, The Unfolded Protein Response and Cell Fate Control. Mol Cell, 2017.

20. Sepulveda, D., et al., Interactome Screening Identifies the ER Luminal Chaperone Hsp47 as a Regulator of the Unfolded Protein Response Transducer IRE1α. Molecular Cell, 2018. 69(2): p. 238–252.e7.

21. Walter, P. and D. Ron, The unfolded protein response: from stress pathway to homeostatic regulation. Science, 2011. 334(334): p. 1081–6.

22. Yoshida, H., et al., XBP1 mRNA is induced by ATF6 and spliced by IRE1 in response to ER stress to produce a highly active transcription factor. Cell, 2001. 107(7): p. 881–91.

23. Shen, X., et al., Complementary signaling pathways regulate the unfolded protein response and are required for C. elegans development. Cell, 2001. 107(7): p. 893–903.

24. Liou, H.C., et al., A new member of the leucine zipper class of proteins that binds to the HLA DR alpha promoter. Science, 1990. 247(247): p. 1581–4.

25. Hetz, C., K. Zhang, and R.J. Kaufman, Mechanisms, regulation and functions of the unfolded protein response. Nat Rev Mol Cell Biol, 2020. 21(8): p. 421–438.

26. Maurel, M., et al., Getting RIDD of RNA: IRE1 in cell fate regulation. Trends Biochem Sci, 2014. 39(5): p. 245–54.

27. Liu, Y., et al., Role for the endoplasmic reticulum stress sensor IRE1α in liver regenerative responses. J Hepatol, 2015. 62(3): p. 590–8.

28. Argemí, J., et al., X-box Binding Protein 1 Regulates Unfolded Protein, Acute-Phase, and DNA Damage Responses During Regeneration of Mouse Liver. Gastroenterology, 2017. 152(5): p. 1203–1216.e15.

29. Hur, K.Y., et al., IRE1α activation protects mice against acetaminophen-induced hepatotoxicity. The Journal of Experimental Medicine, 2012. 209(2): p. 307–318.

30. Hollien, J., et al., Regulated Ire1-dependent decay of messenger RNAs in mammalian cells. The Journal of Cell Biology, 2009. 186(3): p. 323.

31. Hollien, J., et al., Regulated Ire1-dependent decay of messenger RNAs in mammalian cells. Journal of Cell Biology, 2009. 186(3): p. 323–331.

32. Kim, R.S., et al., The XBP1 Arm of the Unfolded Protein Response Induces Fibrogenic Activity in Hepatic Stellate Cells Through Autophagy. Sci Rep, 2016. 6: p. 39342.

33. Heindryckx, F., et al., Endoplasmic reticulum stress enhances fibrosis through IRE1α-mediated degradation of miR-150 and XBP-1 splicing. EMBO Molecular Medicine, 2016. 8(7): p. 729–744.

34. Thamsen, M., et al., Small molecule inhibition of IRE1α kinase/RNase has anti-fibrotic effects in the lung. PLoS One, 2019. 14(1): p. e0209824.

35. Sato, H., et al., 4μ8C Inhibits Insulin Secretion Independent of IRE1α RNase Activity. Cell Struct Funct, 2017. 42(1): p. 61–70.

36. Rufo, N., et al., Stress-induced inflammation evoked by immunogenic cell death is blunted by the IRE1α kinase inhibitor KIRA6 through HSP60 targeting. Cell Death & Differentiation, 2022. 29(1): p. 230–245.

37. Harnoss, J.M., et al., Disruption of IRE1α through its kinase domain attenuates multiple myeloma. Proceedings of the National Academy of Sciences, 2019. 116(33): p. 16420–16429.

38. Lebeaupin, C., et al., Bax inhibitor-1 protects from nonalcoholic steatohepatitis by limiting inositol-requiring enzyme 1 alpha signaling in mice. Hepatology, 2018. 68(2): p. 515–532.

39. Lee, A.H., et al., Regulation of hepatic lipogenesis by the transcription factor XBP1. Science, 2008. 320(320): p. 1492–6.

40. Wang, S., et al., IRE1α-XBP1s induces PDI expression to increase MTP activity for hepatic VLDL assembly and lipid homeostasis. Cell metabolism, 2012. 16(4): p. 473–486.

41. Kühn, R., et al., Inducible gene targeting in mice. Science, 1995. 269(269): p. 1427–9.

42. Iwawaki, T., et al., Function of IRE1 alpha in the placenta is essential for placental development and embryonic viability. Proceedings of the National Academy of Sciences, 2009. 106(39): p. 16657–16662.

43. Campos, G., et al., The transcription factor CHOP, a central component of the transcriptional regulatory network induced upon CCl4 intoxication in mouse liver, is not a critical mediator of hepatotoxicity. Arch Toxicol, 2014. 88(6): p. 1267–80.

44. Lee, A.H., N.N. Iwakoshi, and L.H. Glimcher, XBP-1 regulates a subset of endoplasmic reticulum resident chaperone genes in the unfolded protein response. Mol Cell Biol, 2003. 23(21): p. 7448–59.

45. Jara, C., et al., Genetic ablation of tau improves mitochondrial function and cognitive abilities in the hippocampus. Redox Biol, 2018. 18: p. 279–294.

46. Brown, M.R., P.G. Sullivan, and J.W. Geddes, Synaptic Mitochondria Are More Susceptible to Ca2+Overload than Nonsynaptic Mitochondria*. Journal of Biological Chemistry, 2006. 281(17): p. 11658–11668.

47. Weber, L.W., M. Boll, and A. Stampfl, Hepatotoxicity and mechanism of action of haloalkanes: carbon tetrachloride as a toxicological model. Crit Rev Toxicol, 2003. 33(2): p. 105–36.

48. Zhang, K., et al., The unfolded protein response transducer IRE1α prevents ER stress-induced hepatic steatosis. Embo j, 2011. 30(7): p. 1357–75.

49. Wong, R.J., et al., Nonalcoholic steatohepatitis is the second leading etiology of liver disease among adults awaiting liver transplantation in the United States. Gastroenterology, 2015. 148(3): p. 547–55.

50. Diehl, A.M. and C. Day, Cause, Pathogenesis, and Treatment of Nonalcoholic Steatohepatitis. N Engl J Med, 2017. 377(21): p. 2063–2072.

51. Schwabe, R.F., I. Tabas, and U.B. Pajvani, Mechanisms of Fibrosis Development in Nonalcoholic Steatohepatitis. Gastroenterology, 2020. 158(7): p. 1913–1928.

52. Zito, E., et al., Endoplasmic Reticulum Thiol Oxidase Deficiency Leads to Ascorbic Acid Depletion and Noncanonical Scurvy in Mice. Molecular Cell, 2012. 48(1): p. 39–51.

53. Bickel, M., et al., Selective inhibition of hepatic collagen accumulation in experimental liver fibrosis in rats by a new prolyl 4-hydroxylase inhibitor. Hepatology, 1998. 28(2): p. 404–11.

54. Baumann, S. and T. Hennet, Collagen Accumulation in Osteosarcoma Cells lacking GLT25D1 Collagen Galactosyltransferase. J Biol Chem, 2016. 291(35): p. 18514–24.

55. Lekszas, C., et al., Biallelic TANGO1 mutations cause a novel syndromal disease due to hampered cellular collagen secretion. Elife, 2020. 9.

56. Ishida, Y., et al., Autophagic elimination of misfolded procollagen aggregates in the endoplasmic reticulum as a means of cell protection. Molecular biology of the cell, 2009. 20(11): p. 2744–2754.

57. Peterkofsky, B., Ascorbate requirement for hydroxylation and secretion of procollagen: relationship to inhibition of collagen synthesis in scurvy. Am J Clin Nutr, 1991. 54(6 Suppl): p. 1135s-1140s.

58. Kao, W.W.-Y., J.G. Flaks, and D.J. Prockop, Primary and secondary effects of ascorbate on procollagen synthesis and protein synthesis by primary cultures of tendon fibroblasts. Archives of Biochemistry and Biophysics, 1976. 173(2): p. 638–648.

59. Schwarz, R.I., Procollagen secretion meets the minimum requirements for the rate-controlling step in the ascorbate induction of procollagen synthesis. Journal of Biological Chemistry, 1985. 260(5): p. 3045–3049.

60. Peterkofsky, B., The effect of ascorbic acid on collagen polypeptide synthesis and proline hydroxylation during the growth of cultured fibroblasts. Archives of Biochemistry and Biophysics, 1972. 152(1): p. 318–328.

61. Ellgaard, L. and L.W. Ruddock, The human protein disulphide isomerase family: substrate interactions and functional properties. EMBO Rep, 2005. 6(1): p. 28–32.

62. John, D.C., M.E. Grant, and N.J. Bulleid, Cell-free synthesis and assembly of prolyl 4-hydroxylase: the role of the beta-subunit (PDI) in preventing misfolding and aggregation of the alpha-subunit. Embo j, 1993. 12(4): p. 1587–95.

63. Sheng, X., et al., IRE1α-XBP1s pathway promotes prostate cancer by activating c-MYC signaling. Nature Communications, 2019. 10(1): p. 323.

64. Lhomond, S., et al., Dual IRE1 RNase functions dictate glioblastoma development. EMBO Mol Med, 2023. 15(2): p. e16731.

65. Govaere, O., et al., Transcriptomic profiling across the nonalcoholic fatty liver disease spectrum reveals gene signatures for steatohepatitis and fibrosis. Science Translational Medicine, 2020. 12(572): p. eaba4448.

66. Govaere, O., et al., A proteo-transcriptomic map of non-alcoholic fatty liver disease signatures. Nature Metabolism, 2023.

67. Acosta-Alvear, D., et al., XBP1 controls diverse cell type- and condition-specific transcriptional regulatory networks. Mol Cell, 2007. 27(1): p. 53–66.

68. Zhan, Z., et al., Identification of key genes, pathways and potential therapeutic agents for liver fibrosis using an integrated bioinformatics analysis. PeerJ, 2019. 7: p. e6645.

69. Shoulders, M.D. and R.T. Raines, Collagen structure and stability. Annual review of biochemistry, 2009. 78: p. 929–958.

70. El-Gazzar, A., et al., Bi-allelic mutation in SEC16B alters collagen trafficking and increases ER stress. EMBO Mol Med, 2023: p. e16834.

71. Myllyharju, J. and K.I. Kivirikko, Collagens, modifying enzymes and their mutations in humans, flies and worms. Trends Genet, 2004. 20(1): p. 33–43.

72. Myllyharju, J., Prolyl 4-hydroxylases, the key enzymes of collagen biosynthesis. Matrix Biol, 2003. 22(1): p. 15–24.

73. Kivirikko, K.I. and T. Pihlajaniemi, Collagen hydroxylases and the protein disulfide isomerase subunit of prolyl 4-hydroxylases. Adv Enzymol Relat Areas Mol Biol, 1998. 72: p. 325–98.

74. Bulleid, N.J., Disulfide bond formation in the mammalian endoplasmic reticulum. Cold Spring Harb Perspect Biol, 2012. 4(11).

75. Medinas, D.B., P. Rozas, and C. Hetz, Critical roles of protein disulfide isomerases in balancing proteostasis in the nervous system. J Biol Chem, 2022. 298(7): p. 102087.

76. Kadler, K.E., et al., Collagens at a glance. Journal of Cell Science, 2007. 120(12): p. 1955–1958.

77. Wang, Q., et al., Role of XBP1 in regulating the progression of non-alcoholic steatohepatitis. J Hepatol, 2022. 77(2): p. 312–325.

78. Lebeaupin, C., et al., Endoplasmic reticulum stress signalling and the pathogenesis of non-alcoholic fatty liver disease. J Hepatol, 2018. 69(4): p. 927–947.

79. Kohli, E., et al., Endoplasmic Reticulum Chaperones in Viral Infection: Therapeutic Perspectives. Microbiol Mol Biol Rev, 2021. 85(4): p. e0003521.

80. Tarnutzer, K., et al., Collagen constitutes about twelve percent in females and seventeen percent in males of the total protein in mice. bioRxiv, 2022: p. 2022.11.21.517313.

81. Wu, Y., et al., Dynamically remodeled hepatic extracellular matrix predicts prognosis of early-stage cirrhosis. Cell Death & Disease, 2021. 12(2): p. 163.

82. Rojas-Rivera, D., et al., ER stress sensing mechanism: Putting off the brake on UPR transducers. Oncotarget, 2018. 9(28): p. 19461–19462.

83. Rojkind, M., M.-A. Giambrone, and L. Biempica, Collagen Types in Normal and Cirrhotic Liver. Gastroenterology, 1979. 76(4): p. 710–719.

84. Todd, D.J., et al., XBP1 governs late events in plasma cell differentiation and is not required for antigen-specific memory B cell development. Journal of Experimental Medicine, 2009. 206(10): p. 2151–2159.

85. Cubillos-Ruiz, J.R., et al., ER Stress Sensor XBP1 Controls Anti-tumor Immunity by Disrupting Dendritic Cell Homeostasis. Cell, 2015. 161(7): p. 1527–1538.

86. Hu, C.-C.A., et al., XBP-1 regulates signal transduction, transcription factors and bone marrow colonization in B cells. The EMBO journal, 2009. 28(11): p. 1624–1636.

87. Huh, W.J., et al., XBP1 controls maturation of gastric zymogenic cells by induction of MIST1 and expansion of the rough endoplasmic reticulum. Gastroenterology, 2010. 139(6): p. 2038–49.

88. So, J.S., et al., Silencing of lipid metabolism genes through IRE1α-mediated mRNA decay lowers plasma lipids in mice. Cell Metab, 2012. 16(4): p. 487–99.

89. Olivares, S. and A.S. Henkel, The role of X-box binding protein 1 in the hepatic response to refeeding in mice. Journal of Lipid Research, 2019. 60(2): p. 353–359.

90. Hetz, C. and L.H. Glimcher, Fine-tuning of the unfolded protein response: Assembling the IRE1alpha interactome. Mol Cell, 2009. 35(5): p. 551–61.

91. Yu, J., et al., Phosphorylation switches protein disulfide isomerase activity to maintain proteostasis and attenuate ER stress. Embo j, 2020. 39(10): p. e103841.

92. Ishikawa, T., et al., UPR transducer BBF2H7 allows export of type II collagen in a cargo- and developmental stage–specific manner. Journal of Cell Biology, 2017. 216(6): p. 1761–1774.

93. Carreras-Sureda, A., et al., Non-canonical function of IRE1α determines mitochondria-associated endoplasmic reticulum composition to control calcium transfer and bioenergetics. Nature Cell Biology, 2019. 21(6): p. 755–767.

94. McCaughey, J., et al., ER-to-Golgi trafficking of procollagen in the absence of large carriers. J Cell Biol, 2019. 218(3): p. 929–948.

95. Rodriguez, D.A., et al., BH3-only proteins are part of a regulatory network that control the sustained signalling of the unfolded protein response sensor IRE1α. The EMBO Journal, 2012. 31(10): p. 2322–2335.

96. Urra, H., et al., IRE1α governs cytoskeleton remodelling and cell migration through a direct interaction with filamin A. Nat Cell Biol, 2018. 20(8): p. 942–953.

97. Campos, G., et al., Inflammation-associated suppression of metabolic gene networks in acute and chronic liver disease. Arch Toxicol, 2020. 94(1): p. 205–217.

98. Liu, Z., et al., Transforming growth factor β (TGFβ) cross-talk with the unfolded protein response is critical for hepatic stellate cell activation. J Biol Chem, 2019. 294(9): p. 3137–3151.

99. Mederacke, I., et al., High-yield and high-purity isolation of hepatic stellate cells from normal and fibrotic mouse livers. Nat. Protocols, 2015. 10(2): p. 305–315.

100. Cavallaro, M., et al., Impaired generation of mature neurons by neural stem cells from hypomorphic Sox2 mutants. Development, 2008. 135(3): p. 541–557.

101. Schneider, C.A., W.S. Rasband, and K.W. Eliceiri, NIH Image to ImageJ: 25 years of image analysis. Nat Methods, 2012. 9(7): p. 671–5.

102. Carleton’s Histological Technique. Ulster Med J. 1967 Summer;36(2):172.

103. Junqueira, L.C., G. Bignolas, and R.R. Brentani, Picrosirius staining plus polarization microscopy, a specific method for collagen detection in tissue sections. Histochem J, 1979. 11(4): p. 447–55.

104. Rieppo, L., et al., Histochemical quantification of collagen content in articular cartilage. PloS one, 2019. 14(11): p. e0224839–e0224839.

105. Zhao, P., et al., An AMPK-caspase-6 axis controls liver damage in nonalcoholic steatohepatitis. Science, 2020. 367(367): p. 652–660.

106. Rozas, P., et al., Protein disulfide isomerase ERp57 protects early muscle denervation in experimental ALS. Acta Neuropathologica Communications, 2021. 9(1): p. 21.

107. Plate, L., et al., Small molecule proteostasis regulators that reprogram the ER to reduce extracellular protein aggregation. Elife, 2016. 5.

108. Ryno, L.M., et al., Characterizing the Altered Cellular Proteome Induced by the Stress-Independent Activation of Heat Shock Factor 1. ACS Chemical Biology, 2014. 9(6): p. 1273–1283.

109. Benjamini, Y., A.M. Krieger, and D. Yekutieli, Adaptive linear step-up procedures that control the false discovery rate. Biometrika, 2006. 93(3): p. 491–507.

110. Chen, E.Y., et al., Enrichr: interactive and collaborative HTML5 gene list enrichment analysis tool. BMC Bioinformatics, 2013. 14(1): p. 128.

111. Kuleshov, M.V., et al., Enrichr: a comprehensive gene set enrichment analysis web server 2016 update. Nucleic Acids Res, 2016. 44(W1): p. W90–7.

112. Köhler, S., et al., The Human Phenotype Ontology project: linking molecular biology and disease through phenotype data. Nucleic Acids Res, 2014. 42(Database issue): p. D966-74.

113. Ashburner, M., et al., Gene Ontology: tool for the unification of biology. Nature Genetics, 2000. 25(1): p. 25–29.

114. Szklarczyk, D., et al., STRING v10: protein-protein interaction networks, integrated over the tree of life. Nucleic Acids Res, 2015. 43(Database issue): p. D447-52.

115. Xia, J. and D.S. Wishart, Web-based inference of biological patterns, functions and pathways from metabolomic data using MetaboAnalyst. Nature Protocols, 2011. 6(6): p. 743–760.

116. Pang, Z., et al., Using MetaboAnalyst 5.0 for LC–HRMS spectra processing, multi-omics integration and covariate adjustment of global metabolomics data. Nature Protocols, 2022. 17(8): p. 1735–1761.

117. Kruskal, W.H. and W.A. Wallis, Use of Ranks in One-Criterion Variance Analysis. Journal of the American Statistical Association, 1952. 47(260): p. 583–621.

